# Biotic versus environmental controls on microbial degradation of permafrost organic matter

**DOI:** 10.64898/2026.06.03.729924

**Authors:** Rachel Mackelprang, Michael W. Snyder, Samuel E. Barnett, Anne M. Kellerman, Sommer F. Starr, Suzi Arzoumanian, Caroline Maroutian, Jonathan A. Corpeno, Thomas A. Douglas, Ashley Shade, Robert G.M. Spencer

## Abstract

Permafrost thaw exposes ancient organic matter to microbial degradation, which is predicted to release globally significant quantities of greenhouse gases into the atmosphere. Though microorganisms drive these processes, the relative importance of biotic (taxonomic and functional community composition) versus environmental (e.g., soil physicochemistry) drivers and their interactions are unknown. Using a novel *in situ* thaw experiment conducted at the Cold Regions Research and Engineering Laboratory’s Permafrost Tunnel near Fairbanks, Alaska, we experimentally separated the effects of soil physicochemistry and microbial communities under “real-world” thaw conditions. To simulate thaw, active layer soil, Holocene permafrost (2 kya), and Pleistocene permafrost (40 kya) were sterilized, inoculated with microbial communities from the different soils, enclosed in 0.22 µm membrane bags to prevent immigration, and buried in the active layer. We retrieved the bags after two weeks and two months of thaw and characterized microbial community structure (16S rRNA and ITS2 amplicon sequencing), functional potential (metagenome sequencing), and soil organic matter (OM) composition at the molecular level (FT-ICR MS). Soil had a stronger effect on bacterial community and gene assemblages than inoculum, and the effects of inoculum were stronger and longer-lasting on community structure than functional potential. Pleistocene permafrost initially contained approximately eleven times more dissolved organic carbon than the other soils, and was enriched in OM derived from microbial necromass and low molecular weight organic acids. This carbon was rapidly depleted during thaw and OM compositional characteristics became increasingly similar to active layer and Holocene permafrost, paralleling shifts in Pleistocene permafrost functional gene profiles and bacterial community structure towards those of other soils. Overall, this work provides new insights into the susceptibility of OM to microbial degradation in compositionally distinct permafrost soils, and ways in which Pleistocene Yedoma permafrost carbon is likely to be particularly vulnerable to permafrost thaw.

## Introduction

Permafrost soil stores approximately one-quarter to one-half of Earth’s soil carbon, and up to 40% of this permafrost could disappear by the end of the century ^1,2^. Permafrost is edaphically diverse, and differences in physicochemistry can strongly influence thaw outcomes such as the amounts and types of greenhouse gases (GHG) produced when soils thaw ^3,4^. Carbon inputs to late Pleistocene permafrost (formed 120-11.7 thousand years ago (kya)) were largely derived from the grassland-dominated “mammoth steppe” paleoecosystem ^5–10^. Organic matter (OM) deposition, accumulation, and entrainment in permafrost occurred at relatively fast rates that did not permit substantial decomposition, resulting in a biolabile carbon pool ^11–14^. Warmer and wetter conditions during the Holocene (less than 11.7 kya) caused a shift towards shrubby tree-dominated vegetation, slower permafrost formation, and a more weathered stable carbon pool ^15–18^.

Like permafrost itself, permafrost microbial communities vary widely in taxonomic and functional gene structure. These communities reflect the paleoenvironment present at the time of permafrost formation and the conditions experienced through time while entrained in frozen soil ^19–21^. When permafrost thaws, indigenous communities are poised to have first access to newly exposed carbon. However, thaw enables immigration from the active layer, which may introduce taxa that are better adapted to thawed conditions ^20,22^. Whether indigenous communities influence community assembly trajectories and carbon processing during thaw is an unanswered question. Under conditions that favor specialist taxa such as methanogens, the presence or absence of key members can affect carbon turnover and GHG production ^16,23^. On the other hand, community structure may not be as important in permafrost that contains an abundance of labile carbon that can be metabolized by a taxonomically diverse range of organisms ^19^.

Here, we investigated how the combined effects of soil OM composition, permafrost age, and indigenous communities determine thaw trajectories. We hypothesized that: 1) OM is more bioavailable in Pleistocene permafrost than Holocene permafrost and active layer soil. 2) OM composition drives community response to thaw. 3) OM is more important than indigenous microbial communities for community assembly during thaw. Our novel in situ experimental design enabled us to address these questions under real-world conditions and illustrates the value of integrating high-resolution soil carbon and microbial data to disentangle the interacting roles of carbon substrate and microbial communities during permafrost thaw.

## Results

### Experiment overview

We performed thaw experiments using Holocene permafrost (2 kya), Pleistocene permafrost (40 kya) and active layer soil that froze during winter and had not yet undergone spring thaw (Fig. 1). Samples were homogenized, distributed into 0.22 µm nylon membrane bags (preventing immigration from the surrounding soil) and sterilized ^24,25^. Each sterilized sample bag was inoculated with enough active layer soil, Holocene permafrost, Pleistocene permafrost, or a combination of soils to introduce 1×10^6^ bacterial cells. Bags were buried in the active layer at a depth of 45 cm to simulate thaw and retrieved at two time points--two weeks (2w) and two months (2m). Non-sterilized samples that were not thawed are referred to as frozen or T_0_. Soil OM chemistry was characterized at ultrahigh resolution using FT-ICR MS and microbial communities were characterized using 16S rRNA and ITS2 amplicon sequencing and shotgun metagenome sequencing. Archaeal taxa were extremely rare, so we refer to data from 16S rRNA amplicons as bacterial. Metagenomic data are in the form of gene counts derived from annotation against the Carbohydrate-Active enZYmes Glycoside Hydrolase (CAZy GH) database and Kyoto Encyclopedia of Genes and Genomes (KEGG) database.

**Figure 1.**
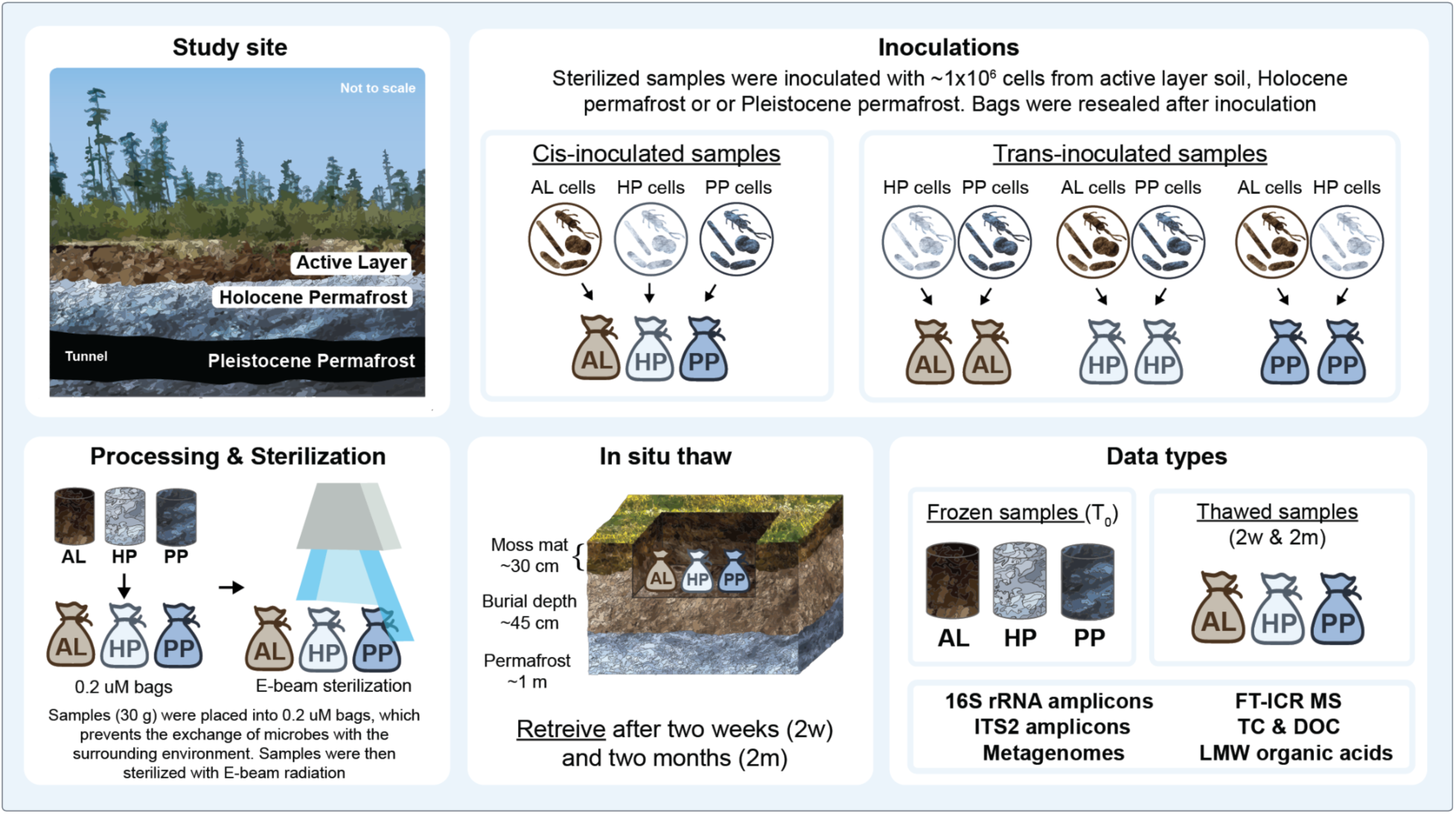
Study design overview. Pleistocene permafrost samples were obtained from within the U.S. Army Cold Regions Research and Engineering Laboratory (CRREL) subsurface Permafrost Tunnel near Fairbanks, Alaska. Active layer and Holocene permafrost were obtained from above the tunnel. Nylon membrane bags (0.22 µm) containing 30 g of homogenized soil were sterilized using E-beam irradiation. Non-sterilized frozen soil was used as the inoculum with the amount adjusted so that approximately 1 × 10^6^ live bacterial cells were added per bag. Samples were divided into categories depending on the inoculum received. Cis-inoculated samples received cells that matched the soil origin (e.g., active layer cells into active layer soil). Trans-inoculated samples received cells that did not match the soil origin (e.g., active layer cells in Pleistocene permafrost). Co-inoculated samples (containing inoculum from two sources) were included in the study but are not shown because they were excluded from analyses that evaluated soil and inoculum effects separately. Bags were buried in late July of 2021 at a depth of 45 cm, which was below the moss mat in silty loess. Samples representing the early stages of thaw were collected after two weeks (2w), and samples representing longer term thaw were collected after two months (2m). LMW: low molecular weight, TC: total carbon.

### Carbon dynamics before and during thaw

Prior to thaw, total carbon (TC) was roughly similar across soil types (2.98 - 3.92%, Table S1), but Pleistocene permafrost was strongly enriched in total nitrogen (TN) relative to the active layer and Holocene permafrost, resulting in significantly lower C:N ratios (one-way ANOVA, Tukey’s HSD, *P <* 0.01, Table S2). DOC was 10.63 and 10.75-fold higher in Pleistocene permafrost than in the active layer and Holocene permafrost, respectively (linear model with Soil x Time interaction, Tukey-adjusted contrasts, *P* < 0.0001, Fig. 2A, Table S3). After two weeks, DOC declined by 74.7% in Pleistocene permafrost (*P* = 4.5 × 10^−6^), 10.9% in Holocene permafrost (not significant), and 13.3% in the active layer (not significant). DOC continued to decline from 2w to 2m in all soils, but the decrease was only significant in the active layer (*P =* 5.0 × 10^−4^). Acetate, butyrate, and propionate pools were found in Pleistocene permafrost before thaw (Table S4). Acetate and butyrate concentrations decreased by >90% within the first two weeks of thaw (Wilcoxon rank-sum test, *P* < 0.01), and propionate decreased by 62% (*P* < 0.05). They were undetectable or found near the limits of quantification in active layer and Holocene permafrost samples.

**Figure 2.**
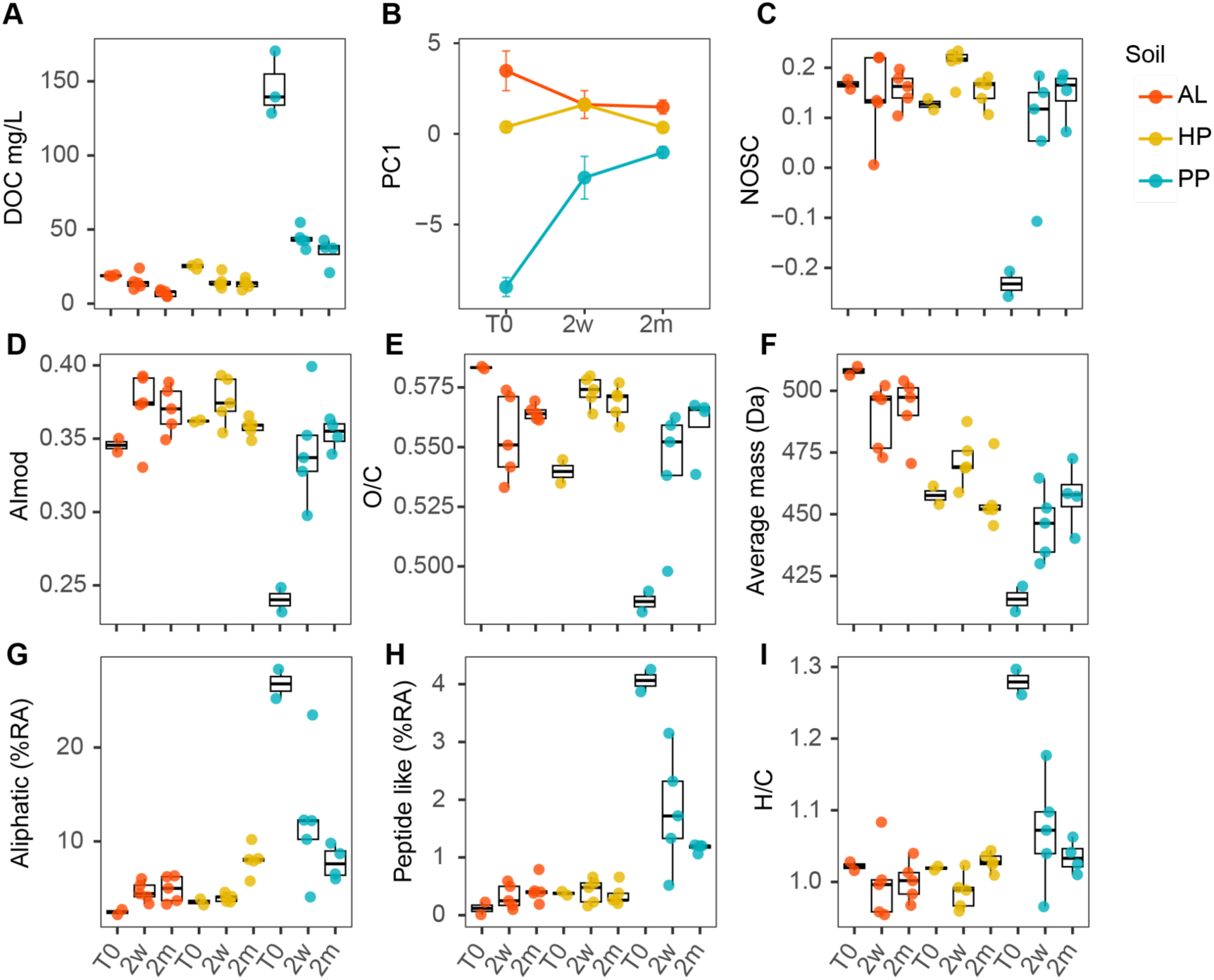
DOM characteristics of active layer soil, Holocene permafrost, and Pleistocene permafrost before thaw (T_0_) and after two weeks (2w) and two months (2m) of thaw. **A)** DOC content. **B)** DOM characteristics represented by plotting the first Principal Component (PC1) scores of FT-ICR MS molecular metrics and compound-class relative abundances. Negative PC1 scores correspond to reduced saturated compound classes (aliphatic, peptide-like, H/C, N/C) that were highest in Pleistocene permafrost at T_0_. Positive PC1 scores correspond to large, oxidized, aromatic formulae (intensity weighted average molecular mass, O/C, NOSC, AImod, polyphenolics, condensed aromatics, and highly unsaturated/phenolic compounds) that were highest in active layer and Holocene permafrost. **C-I)** FT-ICR MS molecular metrics and compound-class relative abundances.

Ultra-high-resolution characterization of molecular formulae from FT-ICR MS showed that composition differed across soils (PERMANOVA, R² = 0.29, pseudo-F = 5.2278, *P* = 0.001), and time points (R² = 0.14, pseudo-F = 2.5703, *P* = 0.0032, Tables S5 & S6). DOC in T_0_ active layer and Holocene permafrost was more oxidized and chemically weathered than in Pleistocene permafrost, as indicated by a higher average molecular mass, greater aromaticity (AImod), higher relative abundance of polyphenolic compounds, and a higher nominal oxidation state of carbon (NOSC) (Fig. 2C-F). In contrast, Pleistocene permafrost DOC was enriched in aliphatic and peptide-like formulae, had higher H/C and N/C ratios, and had a lower NOSC, which is consistent with a reduced, saturated, and biolabile carbon pool derived from microbial sources (e.g., necromass and exopolysaccharides) and non-structural plant polysaccharides (Fig. 2H-I) ^26–28^.

Thaw induced rapid preferential degradation of reduced low molecular weight organic compounds in Pleistocene permafrost, with most of the change occurring in the first two weeks, resulting in a DOC pool that became similar to the active layer and Holocene permafrost (Fig 2B). The relative abundance of aliphatics and peptide-like formulae dropped by 55% and 53% respectively, while average mass, AImod, and polyphenolic relative abundance increased by 7%, 43%, and 75% (Fig 2C-I). DOC composition changed little in the active layer and Holocene permafrost relative to Pleistocene permafrost.

### Interactions between DOC composition, taxa, and genetic potential during thaw

We used network analysis to investigate relationships between DOM composition (FT-ICR MS metrics), microbial communities (bacterial and fungal taxa), and their potential functions (KEGG and CAZy genes) during thaw (Fig. 3). At both the two-week and two-month timepoints, groups of densely connected nodes formed clusters that differentiated Pleistocene permafrost from the active layer and Holocene permafrost (Fig. 3 & S1; Tables S7-S10). The Pleistocene permafrost cluster contained FT-ICR MS metrics indicative of reduced, N-rich, and biolabile OM (H/C, N/C, aliphatics, peptide-like formulae), along with genes and taxa associated with the degradation of microbial necromass, exopolysaccharides, and plant non-structural carbohydrates. FT-ICR MS metrics in the active layer/Holocene permafrost cluster were indicative of stable substrates (polyphenolics, condensed aromatics, O/C, AImod, NOSC, and molecular weight) and contained genes and taxa associated with the degradation of stable substrate, in particular plant cell wall polysaccharides (cellulose, hemicellulose, and pectin).

**Figure 3.**
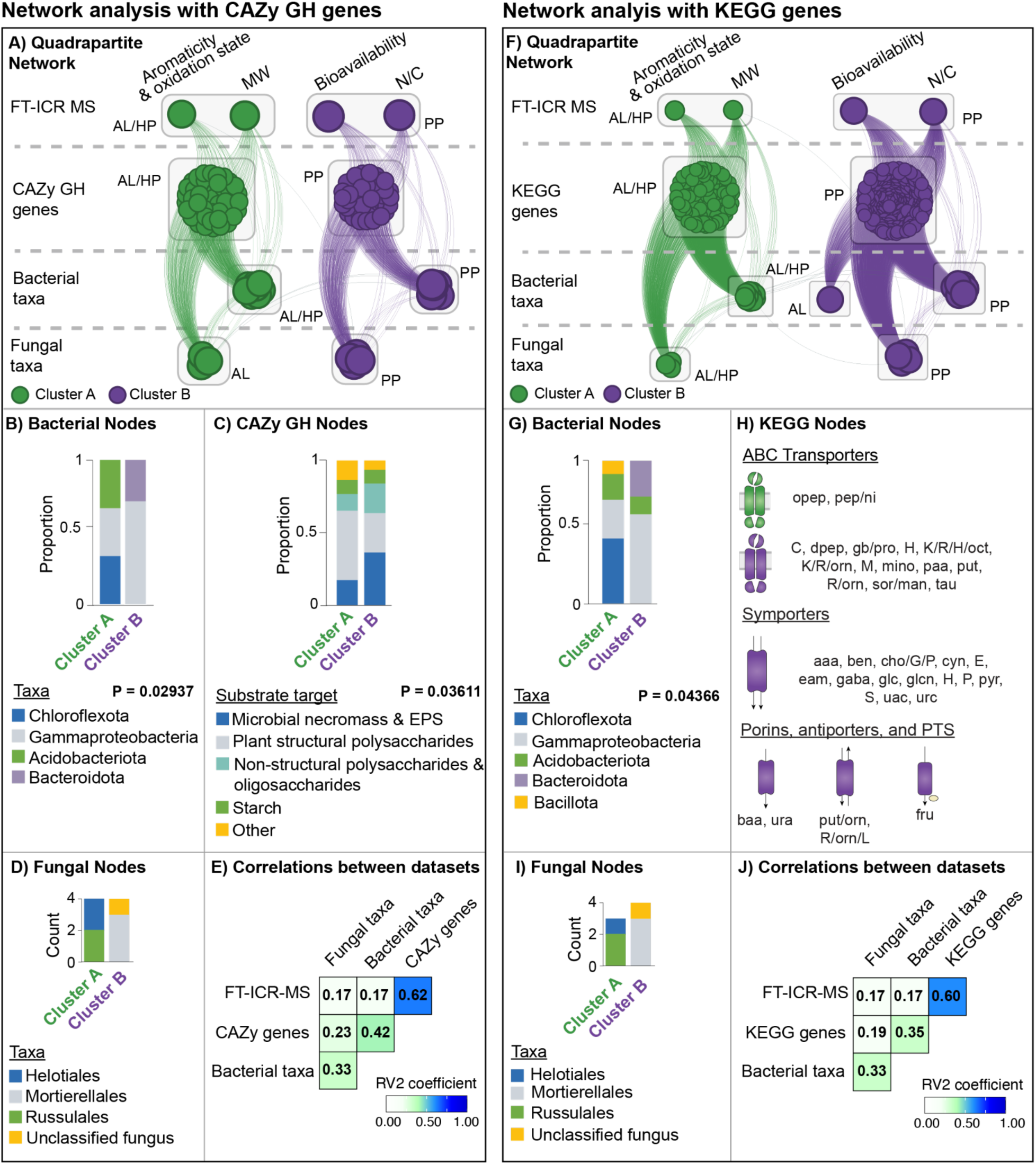
Network analysis of FT-ICR MS, metagenomic, and amplicon data after two weeks of thaw. **A, F.** Network analysis using FT-ICR MS data, CAZy GH or KEGG genes, bacterial ASVs, and fungal ASVs. For FT-ICR MS, nodes represent principal component loadings derived from weighted average molecular metrics and compound class relative abundances. Molecular weight (MW) and N/C were retained as individual variables. The aromaticity and oxidation state node includes polyphenolics, condensed aromatics, NOSC, and O/C. The bioavailability node includes H/C, aliphatics, and peptide-like formulae. Lines show positive correlations between nodes, and node size indicates connectedness. Spin-glass clusters of similarly connected nodes are indicated by color. Semi-transparent boxes indicate the soil in which genes or ASVs were most abundant, or in the FT-ICR-MS metrics were highest. Nodes in a cluster that had different soil abundance profiles than the majority of nodes in that cluster were separated for visualization purposes. **B, G)** Distribution of bacterial taxa within clusters, summarized at the phylum level, except *Gammaproteobacteria*, which is shown at the class level. Fisher’s exact test was used to test whether composition differed among clusters. **D, I)** Counts of fungal ASVs summed at the level of class or order within clusters. **C**) Distribution of CAZy GH genes within clusters grouped by substrate target. Fisher’s exact test was used to test whether substrate groups differed among clusters. **H)** KEGG genes related to the uptake of carbon substrate separated by transporter type and cluster. Green indicates enrichment in Cluster A and purple indicates enrichment in Cluster B. Capital letters are canonical one-letter codes for the twenty standard amino acids. Other abbreviations are opep: oligopeptide, pep: peptide, ni: nickel, dpep: dipeptide, gb: glycine betaine, pro: proline, mino: myo-inositol, paa: polar amino acid, put: putrescine, orn: ornithine, sor: sorbitol, man: mannitol, tau: taurine, baa: basic amino acid, ura: urea, fru: fructose, aaa: aromatic amino acid, aket: alpha-ketoglutarate, ben: benzoate, cho: choline, cyn: cyanate, eam: ethanolamine, glc: glycolate, glcn: gluconate, pyr: pyruvate, uac: uric acid, urc: uracil. **E, J)** Heatmap of RV2 coefficients measuring correlations between data sets.

At two weeks, the potential for polysaccharide degradation differed between clusters (Fisher’s exact test, *P* = 0.036). CAZy GH family genes targeting microbial necromass and exopolysaccharides were more abundant in the Pleistocene permafrost cluster than in the active layer/Holocene permafrost cluster (37% and 17%, respectively) (Fig 3C, Table S7). GH families targeting plant cell wall polysaccharides showed the opposite trend, at 48% in the active layer/Holocene permafrost cluster versus 27% in Pleistocene permafrost.

To determine whether clusters differed in their ability to import carbon substrate, we examined KEGG transport genes (Table S8). Only five transporter genes were found in the 2w active layer/Holocene cluster, compared to 57 in the Pleistocene permafrost cluster. These transporters were consistent with the uptake of substrate derived from microbial necromass, exometabolites, or other labile N-rich pools, including genes for the uptake of: (i) cell components such as N-acetylglucosamine and amino sugars, (ii) cytoplasmic polymers and monomers including peptides, amino acids, polyamines, and osmolytes, (iii) exopolysaccharide-derived substrate, and (iv) small molecules typically found in the exometabolite pool (Fig. 3H).

Taxonomic differences between clusters were consistent with functional gene patterns. Active layer/Holocene permafrost clusters were enriched in *Acidobacteria* and *Chloroflexi* taxa that are oligotrophic degraders of cell-wall polysaccharides in cold soils ^29–32^ (Fisher’s exact test, *P <* 0.05; Fig. 3B, 3G). Bacterial taxa in Pleistocene permafrost clusters were dominated by copiotrophic members of *Gammaproteobacteria*, specifically from the *Pseudomonas* genus and *Oxalobacteraceae* family ^33–35^ (Tables S7 & S8). Fungi in active layer/Holocene clusters include lineages of wood decomposers from the *Leotiomycetes* and *Agaricomycetes* families (Fig 3D, 3I) ^36,37^. Fungi in the Pleistocene permafrost clusters are members of order *Mortierellales* that tend to exploit labile or intermediate carbon pools ^38–41^.

We quantified the strength of associations between datasets using the RV2 coefficient (a multivariate measure of similarity). At two weeks, FT-ICR MS metrics were strongly correlated with KEGG and CAZy genes (RV2 = 0.60 and 0.62, respectively) (Fig. 3E, 3J). Correlations between FT-ICR MS metrics and bacterial taxa were comparatively weak (RV2 = 0.33) and even weaker for fungal taxa (RV2 = 0.17).

### Microbial community assembly

In frozen soils, bacterial taxa and gene abundances (KEGG and CAZy) differed between the active layer, Holocene permafrost, and Pleistocene permafrost (Fig. S2). The most abundant bacterial phyla were *Pseudomonodota, Acidobacteriota*, *Bacillota,* and *Chloroflexota* and the majority of fungal taxa were from the *Helotiales, Thelebolales, and Mortierellales* orders (Fig. S3 & S4).

Thaw shifted community and functional gene composition rapidly, with most change occurring in the first two weeks (Kruskal-Wallis test, *P* < 0.01, Fig. S4). Comparatively little change occurred from two weeks to two months. During thaw, soil, inoculum, and time affected community composition and gene abundances (PERMANOVA, *P <* 0.05, Fig. 4, Table S11). For KEGG and CAZy genes, multivariate dispersion differed among soils, but PERMANOVA results were confirmed using distance-based redundancy analysis (Table S11). Soil was the strongest driver of assembly processes for bacterial communities, KEGG genes, and CAZy genes, explaining 26-52% of the variation in structure versus 4-9% for inoculum. However, inoculum was more important in 16S rRNA amplicon data than functional gene data, explaining 10% of the variation in bacterial communities versus 4% for KEGG and CAZy genes. For fungi, inoculum explained more variation in community structure than soil (15% and 11%, respectively).

**Fig. 4.**
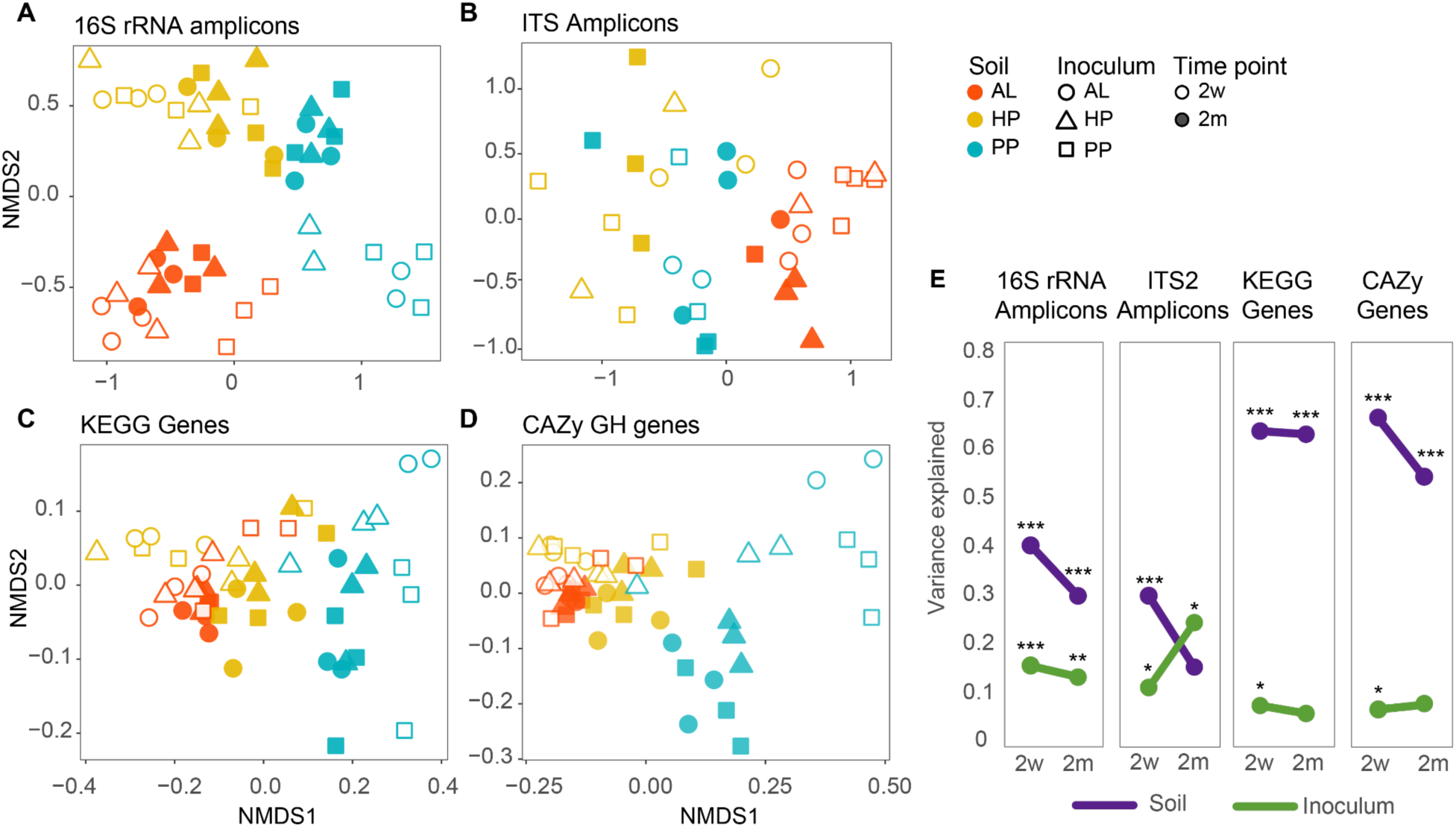
Characteristics of microbial communities and functional genes during thaw. **A-D)** NMDS ordination plots of microbial community and genetic structure. Points are colored by soil, shape shows inoculum, and shape fill indicates timepoint. **E)** Proportion of variance explained by soil (purple) and inoculum (green) during thaw. Data include two-week and two-month timepoints for 16S rRNA amplicons, ITS2 amplicons, KEGG genes, and CAZy GH genes. Variance explained is shown as R² values from PERMANOVA models. Stars indicate significance of each factor on community or genetic structure. ***P < 0.0005, **P < 0.005, *P < 0.05. AL: active layer, HP: Holocene permafrost, PP: Pleistocene permafrost, 2w: two weeks, 2m: two months.

We investigated how long the effects of soil versus inoculum were detectable by comparing the variance explained in PERMANOVA models at two weeks versus two months (Table S12). For KEGG and CAZy genes, multivariate dispersion differed among soils, but PERMANOVA results were confirmed using distance-based redundancy analysis (dbRDA). The effects of inoculum decreased over time for bacterial taxa and functional genes (Fig. 4E). It was no longer significant in the functional gene data after two months but remained significant for bacteria. The effect of soil remained strong at two months, particularly in the KEGG and CAZy gene data, where it explained 63% and 50% of the variation, respectively. The drivers of community assembly were different for fungal communities than the other data types, which showed an increase in the importance of inoculum after two months and a decrease in the variance explained by soil.

We next examined drivers of assembly processes during thaw. We asked if there was evidence of homogenizing selection (driving genes and communities to become more similar over time) or variable selection (driving genes and communities to diverge over time), as indicated by distances of each sample to the centroid of its group. In bacterial taxa, KEGG genes, and CAZy genes, distances to centroids decreased from two weeks to two months (PERMDISP, *P* ≤ 0.023, Fig. S5, Table S13), indicative of homogenizing selection. Fungal community structure did not become more similar during thaw, indicating that homogenizing selection did not play a significant role in structuring fungal communities, at least at the time points observed in this study.

We also investigated assembly processes in bacterial taxa by comparing observed patterns to null models. Specifically, we used the beta nearest taxon index (β-NTI), which measures whether communities are more or less phylogenetically similar than expected by chance ^42^ and Sloan’s neutral modeling framework, which tests whether taxon abundances and distributions can be explained by neutral processes (Fig. S6). We observed evidence of weak to moderate homogenizing selection after two weeks (β-NTI near or below −2). After two months, patterns were more consistent with stochastic assembly processes (β-NTI between −2 and 2). Thaw improved the fit to the Sloan neutral model, which increased from two weeks (R^2^ = 0.563) to two months (R^2^ = 0.618).

### Taxa, genes and pathways differing between soils during thaw

Because environmental selection (i.e., soil) was a key driver of assembly processes during thaw, we identified taxa, genes, and pathways that differed significantly between soils, while controlling for the effects of inoculum. Consistent with network-based analyses, bacteria enriched in the active layer and Holocene permafrost included *Acidobacteriota*, *Chloroflexota, Pseudomonodota*. Within *Pseudomonodota,* active layer and Holocene permafrost enriched ASVs included chemolithotrophic and/or uncharacterized taxa (families *Gallionellaceae*, *Nitrosomonadaceae*, and WD206), as well as the genus *Massilia* (family *Oxalobacteraceae*) ^43–45^. In contrast, ASVs enriched in Pleistocene permafrost were dominated by metabolically versatile and copiotrophic members of *Oxalobacteraceae* (genera *Janthinobacterium* and *Herbaspirillum*) and *Pseudomonadaceae* families (genus *Pseudomonas*) (DESeq2, *P* < 0.05, Tables S14-S20) ^46–49^.

We next identified KEGG genes that were differentially abundant across soils (DESeq2, *P <* 0.05, Fig. S7, Tables S21-S26) and focused on genes related to carbon substrate transport and use (Fig. 5A, Fig. S8, Tables S27-S28). At two weeks, Pleistocene permafrost was enriched in amino acid uptake and degradation genes, particularly for amino acids that are highly abundant in soil (glutamate and aspartate), require substantial ATP/NAD(P)H investment for synthesis (histidine, methionine, and cystine), or are significant sources of sulfur or nitrogen (methionine, cystine, arginine, histidine, lysine, glutamate) ^50–54^. Methionine salvage, lysine degradation, and several arginine degradation pathways were also enriched in Pleistocene permafrost (Fig. S8A). Conversely, thawed active layer and Holocene permafrost were enriched in transporters for the uptake of substrate derived from the degradation of plant structural polysaccharides and genes related to amino acid biosynthesis. After two months, fewer transporters were significantly differentially abundant between the soils and between two weeks and two months the total number of differentially abundant genes decreased by 46%.

**Fig. 5.**
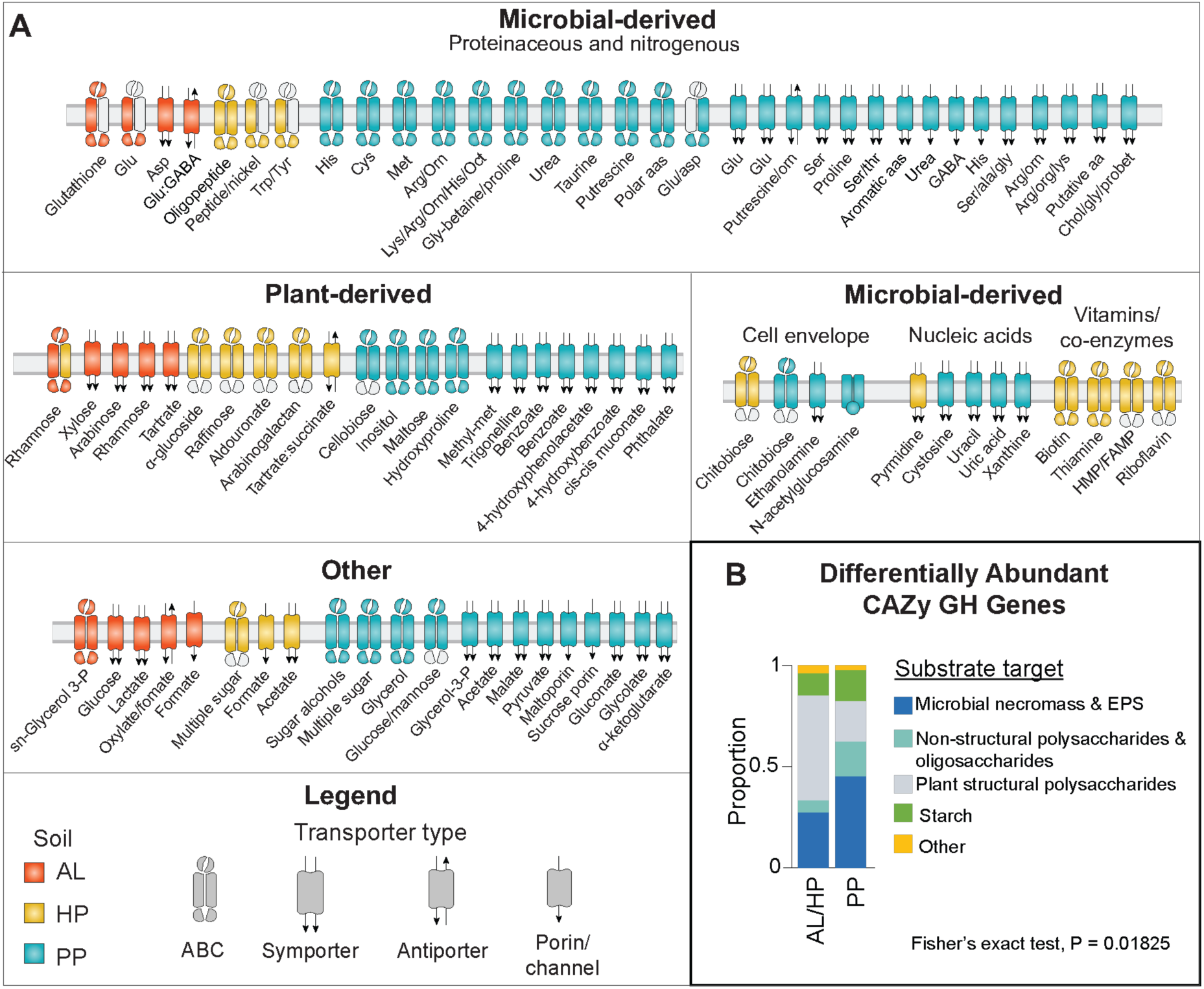
Genes related to carbon and nitrogen substrate uptake and carbohydrate degradation that are differentially abundant across soils after two weeks of thaw. **A)** KEGG transport genes related to the uptake of carbon and nitrogen substrate. Placement into putative microbial-derived or plant-derived categories is based on the likely dominant substrate source during early permafrost thaw. Genes in the “other” category target substrates having more than one potential source or did not clearly fit into the defined categories. The microbial-derived category is further divided into subcategories for proteinaceous or nitrogenous compounds, cell-envelope derived compounds, nucleic acids, and vitamins/co-enzymes. For transporters encoded by multiple genes, colored components represent the genes that were significantly differentially abundant. Amino acids are indicated by canonical three-letter codes. Other abbreviations are defined in Fig 3. KEGG genes represented in the figure and corresponding descriptions are in Table S27. **B)** Differentially abundant CAZy GH families clustered by soil enrichment patterns and grouped by putative substrate target. Fisher’s exact test was used to test whether substrate target groups differed between clusters.

Genes for the degradation of monoaromatic compounds (which are derived from depolymerization of larger aromatic-rich compounds) were also significantly different between soils at the two-week time point (DESeq2, *P <* 0.05) (Fig. S9B). Genes more abundant in the active layer and Holocene permafrost target substituted monoaromatics derived from lignin and humics through protocatechuate and benzoyl-CoA intermediates, while Pleistocene permafrost enriched genes target less-substituted monoaromatics that are processed through the intermediate catechol, which is consistent with a more biolabile carbon pool ^55–57^.

We clustered CAZy GH genes that differed significantly between soils at two weeks (DESeq2, *P <* 0.05, Tables S29-S31), which formed two primary groups differentiating active layer and Holocene permafrost from Pleistocene permafrost (Fig. S10A). Similar to network analysis results, the active layer and Holocene permafrost cluster were enriched in genes that target plant structural polysaccharides (cellulose, hemicellulose, and pectin) and the Pleistocene permafrost cluster had a greater abundance of genes that target microbial-derived substrate (necromass and exopolysaccharides) and non-structural polysaccharides/starch (Fisher’s Exact Test, *P* = 0.01825, Fig. 5B). After two months, the number of genes that differed between soils decreased by 75% (88 to 22) (Fig. S10B, Tables S32-S34).

## Discussion

Microbes both respond to and modify carbon in thawing permafrost. However, we do not know if or how community and functional gene composition affects breakdown, nor how community-soil interactions change across permafrost and soil types that differ by orders of magnitude in age and carbon physicochemistry. Our experimental system enabled us to address these gaps under real-world conditions using manipulations typically limited to the laboratory.

As hypothesized, OM in Pleistocene permafrost was highly biolabile and decomposed quickly when thawed. Carbon from microbial sources, particularly necromass and exopolysaccharides, was a major component of this rapidly disappearing OM pool. Decomposability in Pleistocene permafrost is commonly attributed to rapid permafrost formation (limiting the amount of OM decomposition before entrainment), grass-dominated plant inputs, and pools of low molecular weight organic acids ^13,15,58,59^. Our finding of microbial contributions to Pleistocene OM represents an additional potential major source of biolabile carbon in permafrost.

By directly integrating OM compositional and microbial data, controlling for the effect of taxonomic composition, and investigating functional gene and taxonomic data, we have substantially improved system-level understanding of permafrost thaw. We show that two types of permafrost with similar amounts of total carbon differ substantially in DOM characteristics and thus have large differences in OM turnover and microbial community response to thaw. The depletion of Pleistocene DOC within the first two weeks, combined with the convergence of OM composition, community structure, and functional gene profiles demonstrates that the most reactive fraction of the ancient carbon pool is spent in the earliest stages of thaw and that microbial community and gene composition track the rapid depletion of this biolabile substrate. We also demonstrate that soil characteristics are more important than communities present at the onset of thaw in driving assembly processes, and that community composition has a minimal effect on post-thaw processes.

Permafrost is vast and heterogenous, and the degree to which these patterns generalize will depend on permafrost type and thaw context ^60^. In other conditions, such as carbon-rich permafrost thawed under anaerobic conditions, carbon degradation may be strongly affected by the presence or absence of taxa with phylogenetically constrained metabolisms, such as methanogenesis, where community structure at the onset of thaw may matter more than observed here ^16^. Even within Pleistocene permafrost, variation in paleoenvironment, depositional history, OM composition, and edaphic and landscape conditions would be expected to affect thaw outcomes across sites ^10,20^.

Our results further illustrate that understanding microbial community assembly in thawing permafrost requires both taxonomy and functional genes, and that taxonomy alone risks underestimating or mischaracterizing the environmental drivers of microbial processes. In permafrost and other environments, studies consistently show that environmental factors explain only part of the variation in taxonomic community structure, and a substantial proportion of community variation remains unexplained (e.g., ^20,61–65^). In this study, the strongest microbe-environment links were between OM compositional characteristics and functional genes, whereas correlations with taxa were comparatively weak. In a similar vein, limiting functional genes to specific targets involved in biogeochemical cycles may have missed the cumulative signal from more comprehensive annotations.

These distinctions have important implications for interpreting how microbe-environment interactions affect carbon cycling in thawing permafrost. The apparent mismatch between taxonomic structure and gene distribution can occur in functionally redundant communities, where phylogenetically distinct microorganisms are able to perform similar functions ^66–70^. Environmental filtering in our soils during thaw likely favored metabolic processes, particularly for carbohydrate metabolism and necromass utilization, that are broadly distributed across lineages ^71,72^. The highly abundant taxa we observed in post-thaw samples (*Pseudomonas, Janthinobacterium,* and *Massilia*) are metabolically versatile with phylogenetically dispersed traits, suggesting that generalist functionally redundant taxa are microbial “first-responders” during thaw. This is consistent with previous studies showing that copiotrophic taxa dominate early thaw responses ^35,73–75^.

Together, these results illustrate the value of integrating high-resolution OM chemistry with taxonomic and functional gene data to generate system-level insights. Across analytical approaches, the same soil-level contrasts emerged. The agreement among FT-ICR MS molecular composition, functional gene profiles, transporter genes, CAZy substrate categories, and taxonomic composition indicates that the observed patterns were not specific to a statistical framework. Instead, they point to a coherent coupling between OM composition and microbial functional potential during early thaw. They also indicate that accurate predictions of thaw processes require explicit consideration of soil physicochemistry (including depositional history), and that reliance on taxonomic composition alone is not sufficient to capture the effects of environmental drivers on microbial communities. In many permafrost systems, soil physicochemistry is likely to constrain both community assembly and carbon processing to a far greater degree than taxonomy. Recognizing this hierarchy will be important for interpreting findings across permafrost environments and placing individual studies within a broader framework. Moving forward, comparative work that spans an increasing diversity of permafrost systems and thaw scenarios will move our understanding of permafrost thaw beyond site-specific descriptions and toward generalizable principles that govern thaw outcomes.

## Methods

### Site description and sample collection

Experiments were conducted at the U.S. Army CRREL subsurface Permafrost Tunnel near Fairbanks, Alaska (64.951°N, 147.621°W). The tunnel provides access to Yedoma permafrost, which is silty, organic-rich, syngenetic permafrost that formed in the cold dry mammoth steppe ecosystem of the late Pleistocene ^76,77^. Active layer soil and near-surface permafrost above the tunnel formed during the Holocene, after spruce forests replaced the steppe ecosystem ^78^. Modern vegetation at the site is dominated by black spruce (*Picea mariana*) with a thick cover of sphagnum moss (*Sphagnum spp.*) ^79^. Permafrost at this location and regionally across interior Alaska is actively degrading, characterized by downward expansion of the active layer into thawing near-surface permafrost and widespread emergence of thermokarst pits and erosional gullies ^80,81^.

We previously performed radiocarbon dating of soil 80 m from the tunnel portal, which indicated a permafrost age of 33 kya ^21^. Since then, the tunnel has been expanded and the floor was dropped. Samples for this study were collected 3.5 m below the dated section. Assuming a sediment deposition rate of 0.5 mm per year ^82^, we estimated the approximate age of Pleistocene permafrost used in this study at 40 kya. The age of near surface Holocene permafrost at the depth we sampled (1.5 - 2 m) is estimated to be roughly 2-3 kya based on previous dating and sediment deposition rates ^76,82,83^.

Active layer and Holocene permafrost cores were obtained from above the tunnel by dry drilling to a depth of 2 m using a gasoline-powered 8-cm diameter 1.2 m long Snow, Ice, and Permafrost Research Establishment (SIPRE) corer. Permafrost begins at a depth of 1 m, so for Holocene permafrost we used material from 1.5 - 2 m, which is well below the permafrost table and thus excludes soil susceptible to occasional thaw during warm years. In May 2021, when soil cores were collected, the seasonal thaw layer had expanded downward to a depth of 35 cm. Active layer soil used for this experiment was from 45-80 cm, ensuring that it had not yet experienced summer thaw but was well above the permafrost table. Pleistocene permafrost was obtained from inside the tunnel, roughly 30 m below the surface and 80 m from the portal entrance. Cores were extracted with a SIPRE corer, drilled at a diagonal where the tunnel wall meets the floor. The first 20 cm of each core was discarded to prevent potential effects of being adjacent to the tunnel wall and floor.

Cores were wrapped in aluminum foil and butcher paper, placed into Whirl-Pak bags (Whirl-Pak Filtration Group, Pleasant Prairie, WI), and shipped overnight in coolers. Dry ice lined the top, bottom, and sides of the cooler, followed by a layer of blue ice and fabric. Cores were packed at the center to avoid direct contact with dry ice or blue ice. Once in the lab, cores were stored at −8°C rather than −20°C until processed to more accurately reflect in situ permafrost temperatures at the tunnel site, which are maintained at or below −4°C year round ^84^.

### Core subsectioning

Cores were subsectioned in a room dedicated to processing permafrost samples. Individuals processing the samples wore N95 masks, autoclaved labcoats, hair coverings, sterile nitrile gloves, and goggles. The outer surface of each core (∼2 cm) was removed using a wet tile saw, which was fitted with 25.4 cm autoclaved continuous-rim diamond blades and run without water. The saw stage was covered with sterile aluminum foil. Saw blades, foil, and gloves were replaced frequently.

Subsectioned samples were placed into sterile 50.8 x 38 cm triple-layered Whirl-Pak bags (Whirl-Pak Filtration Group, Pleasant Prairie, WI). 3.1 L autoclavable polypropylene pans were placed on the benchtop and blue ice packs were placed in the bottom of the trays. A sterile towel was layered onto the ice packs, followed by the triple-bagged samples. Another sterile towel was placed over the bag. Samples were homogenized using a rubber mallet, until pieces were roughly the size of a pea (∼1 cm diameter) or smaller. During homogenization, samples were carefully monitored to ensure that bags remained intact and soils remained frozen. Homogenized soil from each layer was set aside, which was used to characterize microbial communities and carbon chemistry prior to thaw. These are referred to as frozen or T_0_ samples.

### Cell enumeration

For enumeration, cells were separated from the soil matrix using Nycodenz density centrifugation and subjected to live/dead staining ^85–88^. Five ml of 2% sodium hexametaphosphate (SHMP) was added to 1 g of still-frozen soil, vortexed briefly, and rotated at 80 rpm for 20 min on a HulaMixer (ThermoFisher) at 4°C. Samples were sonicated on ice for 1 minute with a 0.3 cm probe (QSonica Q125) at 2.5 w and centrifuged for 10 min at 750 x g at 4°C to pellet debris, after which the supernatant was collected. Fresh 2% SHMP was added to the pellet and the process was repeated two additional times. The pooled supernatant was vacuum filtered through sterile gauze to remove large remaining particles. A 1:1 volume of supernatant was layered over a 50% (w/v) Nycodenz density cushion and centrifuged at 14,000 x g for 1 hr at 4°C. The upper layer containing cells was pelleted by centrifugation at 10,000 x g for 15 min at 4°C. Cells were resuspended in 0.85% NaCl.

Dual staining of live and dead cells was performed using the LIVE/DEAD BacLight viability kit per manufacturer instructions (Invitrogen Technologies). Fluorescently stained cells were vacuum filtered onto a 25 mm diameter 0.22 µm black polycarbonate membrane. The membrane was transferred onto a microscope slide and observed at 1000x total magnification on one focal plane using a Zeiss Axio Imager M2 Fluorescence microscope coupled with an Apotome 2.0 system. Fifteen fields of view were counted per soil type and the average number of cells per field of view was multiplied by the area of the filter membrane.

### Sample bag preparation

Nylon membrane bags, which were 10 x 15 cm in size, were prepared by cutting 10 x 30 cm strips of 0.22 µm nylon membrane roll stock (#RS10233, Tisch Scientific,), folding it in half, and sealing the sides with Flex Seal rubberized waterproof tape. 30 g of homogenized sample was distributed into the bags, and the top was sealed with waterproof tape. Soil bags were sterilized at Steri-Tek (Fremont, CA) using 27 kGy electron-beam (E-beam) irradiation, which potentially causes less alteration of OM than autoclaving or gamma irradiation ^89,90^.

After sterilization, each bag was inoculated with non-sterilized homogenized active layer soil, Holocene permafrost, Pleistocene permafrost, or a combination of two soils. To decouple the effects of community composition from cell abundance, the amount of soil was adjusted so that 1×10^6^ live cells were added per bag. Inoculation was performed by clipping off a small corner of each bag with sterilized scissors, adding the inoculum, and resealing the bag with sterilized Flex Seal tape. Inoculations were organized into three categories, cis-inoculated, trans-inoculated, and co-inoculated. In cis-inoculated treatments, communities were reintroduced into soils from the same origin (e.g., Pleistocene permafrost communities into Pleistocene permafrost). In trans-inoculated treatments, soils and communities originated from different sources (e.g., active layer communities into Pleistocene permafrost). In co-inoculated treatments, mixed communities from two sources were inoculated into one soil (e.g., active layer communities and Pleistocene permafrost communities into Pleistocene permafrost). Three replicate bags were prepared for each soil-community-timepoint combination. We note that co-inoculated samples were used only for analyses where inoculum was not included as a parameter in the statistical model because the potential effects of co-inoculation are not well understood and thus could not accurately be represented.

### Sample burial and retrieval

Samples were buried in late July 2021 in the active layer above the tunnel at a depth of 45 cm. This depth was chosen because it is in the loess layer above the permafrost in soils that had thawed to 50 cm at the time of burial. We dug 2×2 m pits by first cutting out and lifting the moss mat and then removing soil to the desired depth. Bags were sandwiched between galvanized steel hardware cloth and landscape fabric, after which the excavated soil and moss mat were replaced. The first batch of samples was retrieved in mid-August, after two weeks of thaw. The second batch was retrieved in late September, after two months of thaw. Samples were shipped in coolers using the same packing technique that was used for the soil cores, as described above.

### Soil chemistry characterization

#### Total carbon, total nitrogen, and low molecular weight organic acids

Total carbon and total nitrogen were measured by dry combustion using a C/N elemental analyzer at Regen Ag Lab. For organic acids, 5 g of frozen soil was combined with 2.5 ml HPLC-grade water, extracted with gentle shaking for 1 hr, and centrifuged at 10000 x g for 15 min. The supernatant was passed through a sterile 0.2 µm polyethersulfone (PES) filter. Filtrates were analyzed by high-performance liquid chromatography using a MetaCarb 67H organic acids column (0.65 x 30 cm) with H_2_SO_4_ eluent at a flow rate of 0.8 mL/min.

#### Soil leachates

Biological replicates were pooled before leaching water-soluble organic matter because individual samples would not yield sufficient DOC for concentration measurements and downstream FT-ICR MS. Multiple pooled samples representing different inoculum treatments within each soil provided independent assessments of OM composition across treatments. First, frozen replicate samples were pooled in muffled glassware (550°C > 5 hr). DOM was leached at a ratio of 1:5 soil to ultrapure water by weight. Samples were shaken to incorporate and stored in the dark at room temperature for 24 hours. The leachate was then filtered through pre-combusted 0.7 µm glass fiber filters (450°C > 5 hr). Aliquots for DOC concentration and solid phase extraction were acidified to pH 2 with 12 M analytical grade hydrochloric acid and stored at 4°C in the dark until analysis.

#### Dissolved organic carbon quantification

Filtered, acidified samples were analyzed for DOC concentration using a Shimadzu TOC-L high-temperature catalytic oxidation total organic carbon analyzer. Samples were sparged for 5 minutes with a flow rate of 80 ml min^−1^ before quantification of nonpurgeable organic carbon. Quantification was based on a 6-point calibration curve of potassium hydrogen phthalate, and all replicate injections had a coefficient of variance < 2%.

#### Solid phase extraction and Fourier transform ion cyclotron resonance mass spectrometry

Samples were solid phase extracted as described by Dittmar et al., (2008) with modifications described by Starr et al., 2025 ^91,92^. Cartridges used for solid phase extraction (100 mg PPL cartridges; Agilent Technologies) were conditioned by rinsing and soaking the cartridges overnight with HPLC grade methanol. Cartridges were then rinsed with two cartridge volumes of ultrapure water, then another cartridge volume of methanol, before being primed with pH 2 ultrapure water. Sample aliquots were then passed through the cartridges, the volumes of which were determined based on the DOC concentration to achieve a target extract concentration of 40 µg C L^−1^. Cartridges were then rinsed with 2 cartridge volumes of pH 2 ultrapure water, dried under a stream of N_2_ gas, and then eluted with 1 mL of HPLC grade methanol. DOM extracts were stored at −20°C until analysis.

DOM composition was characterized using electrospray ionization in negative mode on a custom-built hybrid linear ion trap FT-ICR MS housing a 21 T superconducting solenoid magnet ^93,94^. Specific operating conditions for the instrument are described in Starr et al. 2025 ^92^. Acquired mass spectra were phase-corrected, converted to Kendrick mass scale ^95^ and internally calibrated using 10-15 abundant homologous series spanning the entire spectra via walking calibration with PetroOrg © software ^96–98^. Only molecular formulae assignments with an error of fewer than 0.2 parts-per-million were retained for further analysis.

The modified aromaticity index (AImod) ^99^, was calculated for each assigned formula as:

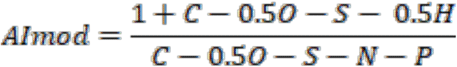

Compound classes were defined based on elemental ratios and the modified aromaticity index (AI_mod_); specifically, low O/C highly unsaturated polyphenolics (HUPs) (AImod < 0.5, H/C < 1.5, O/C < 0.5), high O/C highly unsaturated polyphenolics (HUPs) (AImod < 0.5, H/C < 1.5, O/C ≥ 0.5), aliphatic (H/C ≥ 1.5, N = 0), condensed aromatic (AImod ≥ 0.67), polyphenolic (0.67 > AImod > 0.5), and peptide-like (H/C ≥ 1.5, N > 0). Contributions of each heteroatom and compound class to total composition were calculated as the sum of all the relative abundances (RA) of each peak in a given compound class divided by the total summed abundance of all assigned formulae.

The nominal oxidation state of carbon (NOSC) ^100^, was calculated using the formula:

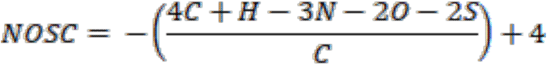

### DNA extraction

Prior to DNA extraction, samples were washed to deplete extracellular DNA using a protocol adapted from ^101,102^. 0.5-0.75 g soil and 800 µl of 4°C sterile phosphate buffer (0.12 M Na2HPO4 [pH 8]) were added to lysing matrix E tubes (MP Biomedicals) and thawed on ice. The slurry was mixed at 300 rpm at 4°C for 15 min and centrifuged at 10,000 x g for 10 min at 4°C. The supernatant was discarded and DNA was extracted from the remaining soil using the FastDNA Spin Kit for Soil (MP Biomedicals). In this protocol, bead beating was conducted for 40 s at 6 m/s. An additional clean-up step to remove humic material and other co-extracted contaminants was performed using the DNeasy Powerclean Pro Cleanup Kit (QIAGEN).

For unincubated T_0_ permafrost samples, DNA purification steps did not sufficiently remove co-extracted contaminants, resulting in failures of downstream PCR and library construction reactions. For these, we re-extracted DNA using the RNeasy PowerSoil Total RNA kit followed by the RNeasy PowerSoil DNA elution kit (QIAGEN), which increased PCR and library construction success rates.

### Amplicon sequencing

We amplified the V4-V5 regions of the 16S rRNA gene using primers 515F and 926R following the Earth Microbiome Project protocol ^103,104^. The fungal ITS2 region was amplified using the 5.8Fun/ITS4-Fun primer pair ^105^. PCR amplification was performed in triplicate 25 µl reactions using recommended reaction and cycling conditions, except that we used Phusion High-Fidelity polymerase (Thermo Scientific) and 1µl of bovine serum albumin. Triplicate PCRs were combined, samples were pooled at equal concentrations, and pooled libraries were purified using the QIAquick PCR Purification Kit (Qiagen). For 16S rRNA amplicons, paired-end (2×250) sequencing was performed on the Illumina MiSeq platform. Because low fungal biomass and co-extracted inhibitors reduced read quality and overlap in some ITS2 libraries, only forward 150-bp reads were retained for ITS2 analyses.

Amplicon data were processed in QIIME2 v2022.2 (for 16S rRNA amplicons) and v2023.7 (ITS2 amplicons) ^106^. Quality filtering, end joining, and amplicon sequence variant (ASVs) generation were performed using the dada2 plugin ^107^. ASVs with fewer than ten reads across all samples were removed prior to downstream analysis. Taxonomic classification of 16S rRNA ASVs was performed using the feature-classifier plugin and the SILVA v138.1 database trained on the 515F-926R region using RESCRIPt ^108,109^. ITS2 taxonomic classification was performed by comparison to the pre-trained dynamic version of the UNITE database (v9.0, 18.07.2023) ^110^. Feature tables, representative sequences, and taxonomic classifications were exported from QIIME2 for downstream statistical analyses in R ^111^.

### Metagenome sequencing and functional annotation

For metagenomic sequencing, library preparation and sequencing was conducted at the Michigan State University Genomics Core. Libraries were prepared using the Kapa HyperPrep Kit (Roche) and paired end sequencing (2×150) was performed on an Illumina NovaSeq 6000 or Element Biosciences AVITI platform. Adapter trimming and quality filtering was performed using bbduk and paired reads were merged using bbmerge (v38.98) ^112,113^.

Reads were annotated by comparison to the Kyoto Encyclopedia of Genes and Genomes (KEGG) (Release 2022-12-26) and Carbohydrate Active EnZYme (CAZy) databases (dbCAN, downloaded 2022-08-06) using DIAMOND (v2.0.15) at an e-value threshold of 1×10^−6 114–116^. Merged reads were assigned the KEGG Orthology (KO) number or CAZy Glycoside Hydrolase (GH) family of the top hit, as described previously ^20,21,117^. Unmerged reads were annotated if the KO number or GH family was the same for both reads. For KEGG and CAZy gene count data, low abundance genes (counts lower than 1000 for KEGG genes and 100 for CAZy genes) were removed. Data were rarefied using the phyloseq package ^118^. Metagenome and amplicon sequence data are available at the NCBI sequence read archive under Bioproject PRJNA1473350.

### Statistical analyses

#### Soil chemistry

All statistical analyses were performed in R ^111^. At T_0_, differences in total carbon (TC), total nitrogen (TN), and C:N among soils were assessed using one-way ANOVA followed by Tukey’s HSD post-hoc test. Prior to analysis, normality was evaluated within each group using the Shapiro-Wilk test and homogeneity of variance was assessed using Levene’s test. Effect sizes for significant pairwise comparisons were calculated as Hedges’ g with 95% confidence intervals using the effectsize package ^119^.

Differences in DOC concentrations were analyzed using a linear model with soil (active layer, Holocene permafrost, Pleistocene permafrost), timepoint (T_0_, two weeks, two months), and their interactions as fixed effects, following log transformation of DOC values. Overall effects were assessed using Type II ANOVA. Pairwise comparisons among soils and timepoints were performed using estimated marginal means with Tukey adjustment implemented in the emmeans package ^120^. Fold differences and percent changes were derived from back-transformed contrasts on the log scale.

Changes in acetate, butyrate, and propionate concentrations in Pleistocene permafrost leachate were assessed using one-sided Wilcoxon rank-sum tests. Normal approximations were used due to some zero values. Active layer and Holocene permafrost samples were excluded because concentrations were at or below the limit of quantification.

FT-ICR MS molecular formula intensity data were Hellinger transformed and Euclidean distances were calculated on the transformed matrix. Differences in molecular composition between soils and time points were tested with permutational multivariate analysis of variance (PERMANOVA) using the adonis2 function in the vegan package ^121^.

FT-ICR MS–derived molecular metrics were summarized for each sample, including intensity-weighted averages (mass, H:C, O:C, N:C, NOSC, and AImod) and relative abundance (%RA) of compound classes. RA variables were treated as compositional data and transformed using a centered log-ratio (CLR) transformation following addition of a small pseudocount (1 × 10⁻⁶) to account for zeros. Weighted-average variables were standardized (z-scored) to unit variance. The transformed compositional variables and scaled weighted metrics were then combined into a single feature matrix and standardized prior to analysis. Principal component analysis (PCA) was performed on the combined dataset (prcomp). Group means and standard errors of PC1 scores were calculated for each soil × timepoint combination and used to visualize trajectories through thaw.

#### Network analysis

We performed quadripartite network analysis by integrating FT-ICR MS summary metrics, functional genes (KEGG genes or CAZy GH gene families), 16S rRNA amplicons, and ITS2 amplicons. Because replicate samples were pooled prior to FT-ICR MS, functional gene and amplicon data replicates were merged network analysis by rarefying counts 100 times (rarefy_even_depth function in the R phyloseq package ^111,118^), averaging counts across iterations, and then averaging counts across replicate samples.

Count-based blocks (genes and amplicon data) were centered log-ratio transformed after addition of a pseudocount of one. For FT-ICR MS metrics, relative abundance variables were scaled by 100 and transformed using log(1+x). Continuous metrics were not transformed. The 12 FT-ICR MS summary metrics exhibited substantial multicollinearity, and Sparse Generalised Canonical Correlation Analysis’ (sGCCA) L1 penalty cannot impose meaningful sparsity on blocks with few variables, so we reduced the FT-ICR MS block prior to multi-block integration. Hierarchical clustering was performed on a signed correlation distance matrix using complete linkage. Signed distances were used rather than absolute value distances so that variables with strong inverse correlations were assigned to separate clusters, preserving the direction of chemical variation for downstream network interpretation. The number of clusters was determined by silhouette analysis. For multi-member clusters, the first principal component (PC1) was used to create a composite variable capturing shared variance. Singleton clusters were retained as scaled single variables. Inter-block similarity was quantified using the RV coefficient in the FactoMineR package ^122^. The RV coefficients were used to construct a weighted design matrix ^123,124^. Block-specific penalties were selected using stability-based tuning separately for each block. The FT-ICR MS block penalty was fixed at 1.0.

We performed unsupervised sGCCA from the mixOmics package ^125,126^. Features associated across blocks were identified by repeated cross-validation. We performed 5-fold cross-validation repeated 200 times (1000 total fits). Within each training fold, amplicon and gene blocks were re-transformed, whereas the reduced FT-ICR MS block was carried forward unchanged. A final sGCCA model was fit using only the retained features for each block, with penalties fixed to 1. The resulting network was extracted with an edge cutoff of 0.5. Nodes in the network were clustered using the cluster_spinglass function from the igraph package ^127^ and networks were visualized in Gephi ^128^. To test whether taxonomic composition or CAZy GH family categories differed among clusters, we used Fisher’s exact test. The strength of associations among gene, amplicon, and FT-ICR MS data was quantified using pairwise RV2 coefficients using the MatrixCorrelation package ^123^. Because our primary goal was to evaluate broad cross-associations between datasets, we interpreted networks at the level of major multiblock clusters and feature categories rather than individual edges, hub nodes, or direct chemistry-gene-taxon interactions.

#### Gene and amplicon data

Relationships among samples were quantified using Bray-Curtis dissimilarities and visualized using nonmetric multidimensional scaling (NMDS) ordination plots within the vegan and ggplot2 packages ^121^. We tested treatment (soil, inoculum, and time) and interaction effects with permutational multivariate analysis of variance (PERMANOVA) using the adonis2 function. Effect sizes for each factor were estimated as the proportion of total variation in the distance matrix explained by the model (R²). We performed multivariate dispersion analysis (PERMDISP) with the betadisper function. To evaluate whether PERMANOVA results for functional gene data were robust to heterogeneity of multivariate dispersion detected by PERMDISP, we performed distance-based redundancy analysis (dbRDA) using the dbrda function in vegan.

We quantified the magnitude of change following thaw using frozen T_0_ and cis-inoculated samples. Bray-Curtis dissimilarities were calculated from square root transformed abundance data. We computed multivariate dispersion (betadisper function), with groups defined by the combination of soil and timepoint. Principal coordinate scores were extracted to define a common ordination space. For each soil, the Euclidean distance between individual two week samples and the centroid of the frozen T_0_ samples was calculated in PCoA space to represent the magnitude of thaw-induced change. Similarly, the distance between two month samples and two week centroids was used to quantify change from two weeks to two months. To account for soil-specific variability, distances were normalized by the mean within-group distance to centroid. Differences in the magnitude of change between T_0_ → two week versus two week → two month were tested using linear models with time interval and soil as fixed effects.

To determine whether communities became more similar through time, we used multivariate dispersion (betadisper package) to calculate the distance of each sample to the centroid of its incubation-time group ^129^. For these analyses, we used cis-, trans-, and co-inoculated samples. Differences in dispersion between the two timepoints (two weeks versus two months) were assessed using a permutation test. Lower average distances to centroids at two months relative to two weeks indicate a decrease in β-diversity and was interpreted as evidence of homogenizing selection.

Community assembly analyses were performed using both the β-nearest taxon index (βNTI) ^130^ and Sloan Neutral Model ^131^ methods. For both methods, separate calculations were performed for each timepoint (*i.e.,* frozen, 2 week, and 2 month). The βNTI method is a null modeling approach to estimate the influence of deterministic processes on community assembly. To calculate βNTI, we measured the observed abundance weighted β-mean-nearest taxon distance (βMNTD) between all samples using function *comdistnt* from the picante package ^132^ then generated a βMNTD null model by randomly shuffling the phylogenetic tree tips and recalculating pairwise βMNTD 999 times. βNTI was then calculated as βNTI = (observed βMNTD - mean of null model βMNTDs) / standard deviation of null model βMNTDs. Only sample pairs that were from the same soil origin were used. Sample pairs with |βNTI| < 2 are expected to result from stochastic processes dominating assembly while sample pairs with βNTI > 2 indicate variable selection and βNTI < −2 indicate homogenizing selection as the dominant assembly processes ^130^. The Sloan Neutral Model states that the probability of a species occurring at a given abundance can be predicted using the beta probability density function. Thus we tested the fit of the neutral model on the relationship between ASV occupancy and abundance across all samples using code modified from Burns et al. 2016 ^133^. Poor model fit (*i.e.,* R^2^) indicates that deterministic processes likely dominate assembly across samples. The model was generated using the function pbeta from package stats and fit to the data using the function *nlsLM* from package <u>minpack.lm</u> with 95% confidence intervals calculated with function *binconf* from package Hmisc.

At each timepoint, we identified genes and taxa that differed between soil types, while controlling for the effects of community source, using a negative binomial model with non-rarefied data, implemented in DESeq2 ^134^. Log2 fold changes and Wald test p-values were computed for each gene across all pairwise soil contrasts (AL vs. HP, AL vs. PP, HP vs. PP). P-values were adjusted using the Benjamini–Hochberg false discovery rate.

We generated clustered heatmaps to visualize patterns of differential gene abundance data among soil origins. Row-scaled abundances were visualized using the pheatmap package ^135^. Counts were z-score transformed across genes and hierarchical clustering of genes was performed using Ward’s minimum variance method. Differentially abundant CAZy GH genes were grouped into categories by primary putative substrate target: 1) plant structural polysaccharides (cellulose, hemicellulose, and pectin), 2) microbial-derived material (necromass and exopolysaccharides), 3) nonstructural polysaccharides, oligosaccharides, and sugars, 4) storage molecules (primarily starch), and 5) other. A Fisher’s exact test was used to test whether GH family categories differed among soil types.

## Acknowledgements

We thank Carolina Gonzalez and Serina Hernandez for help with sampling efforts and John Chodkowski for help with data processing. R.M., and R.G.M.S. were supported by the National Science Foundation Division of Environmental Biology and Office of Polar Programs, grant numbers DEB 2029573 and DEB 2029585 respectively. T.A.D. acknowledges funding from the Department of Defense’s Strategic Environmental Research and Development Program (project RC18-1170), Environmental Science and Technology Certification Program (project NH22-7408), and the Army Research Office’s Laboratory University Collaboration Initiative.

## Author contributions

RM, AS, and RGMS conceived of the study and guided the project. RM, TAD, SA, SFS, CM, JAC, and MWS collected and processed the samples. TAD provided expert information about the field site. RM, MWS, SEB, AMK, and SFS analyzed data. RM and SEB performed statistical analyses. RM wrote the manuscript with input from all authors.

