## Supplemental figures and tables for "Biotic versus environmental controls on microbial degradation of permafrost organic matter"

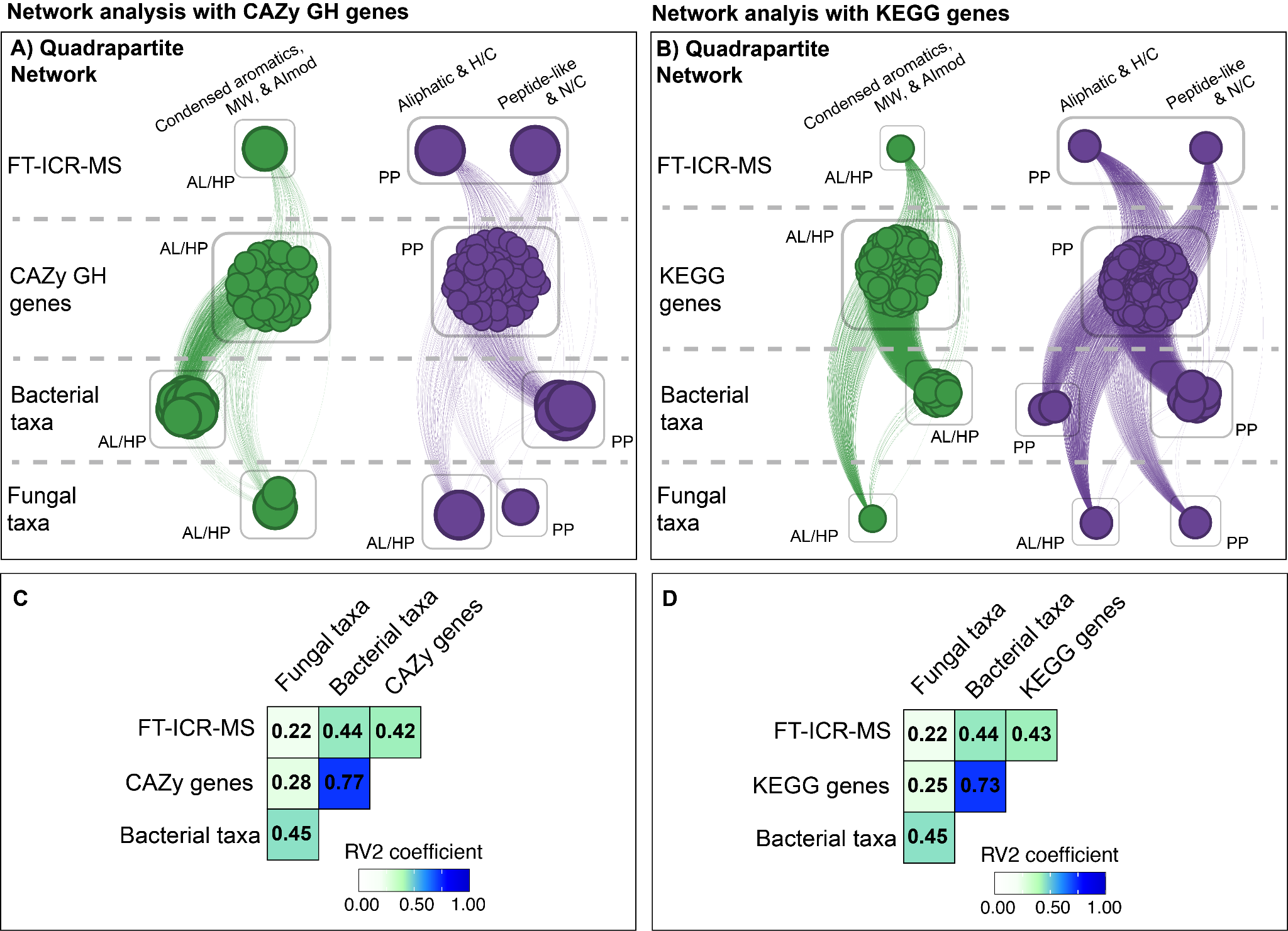

**Figure S1. Network analysis of FT-ICR-MS, metagenomic, and amplicon data after two months of thaw.** Network analysis using FT-ICR-MS metrics, bacterial ASVs, fungal ASVs, and CAZy GH **(A)** or KEGG **(B)** genes. For FT-ICR MS, nodes represent principal component loadings derived from weighted average molecular metrics and compound class relative abundances. Lines show positive correlations between nodes, and node size indicates connectedness. Spin-glass clusters of similarly connected nodes are indicated by color. Soil (AL, HP, or PP) in which genes or ASVs were most abundant or in which FT-ICR-MS metrics had highest value is indicated by semi-transparent boxes drawn around groups of nodes. Nodes in a cluster that had different soil abundance profiles than the majority of nodes in that cluster were manually grouped for visualization purposes. **C, D)** Heatmap of RV2 coefficients measuring correlations between data sets.

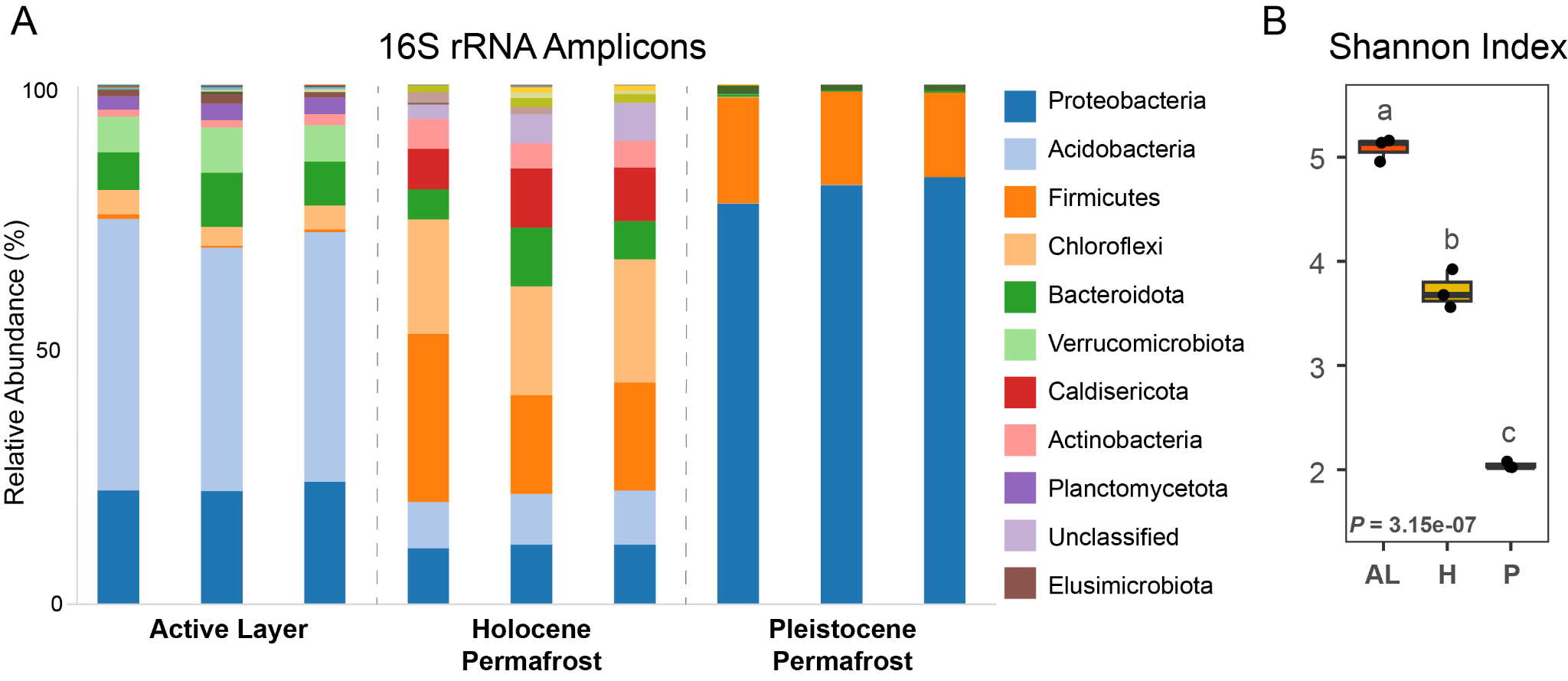

**Supplemental Figure 2. Microbial community structure and alpha diversity in soils prior to thaw.** A) Relative abundance of major phyla from 16S rRNA amplicon data. B) Box plot of the Shannon Diversity Index at the ASV level by soil type (AL: active layer, H: Holocene permafrost, P: Pleistocene permafrost). The *P-*value at the bottom of the graph is the overall effect of soil type on alpha diversity (one-way ANOVA). Different lower-case letters represent statistical differences between soil types (Tukey post-hoc test). In all cases, *P* ≤ 2.8e-05.

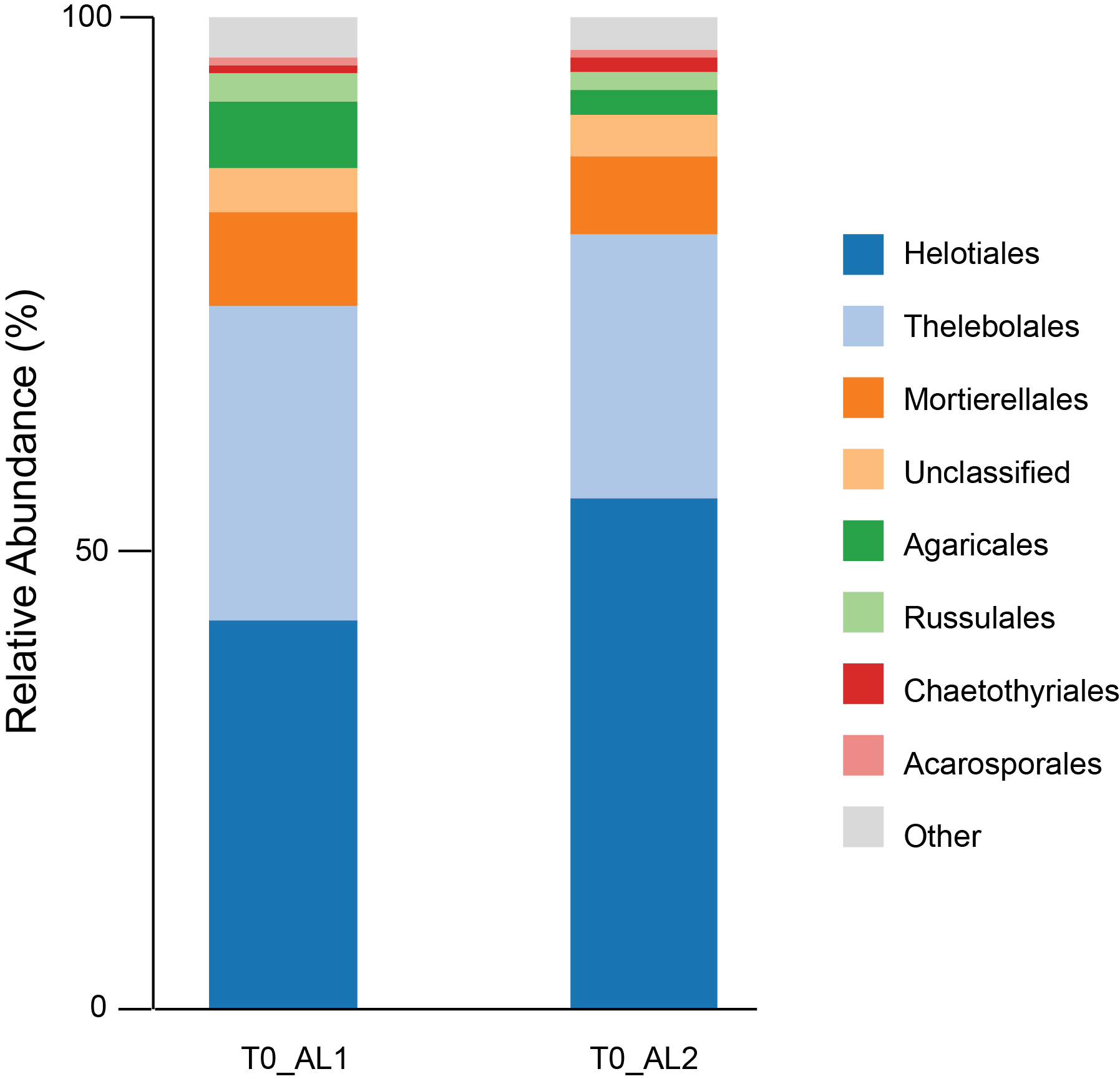

**Supplemental Figure 3. Relative abundance of fungal orders in the active layer prior to thaw.** Only the active layer is shown because amplification of the ITS2 region in T_0_ permafrost samples was unsuccessful.

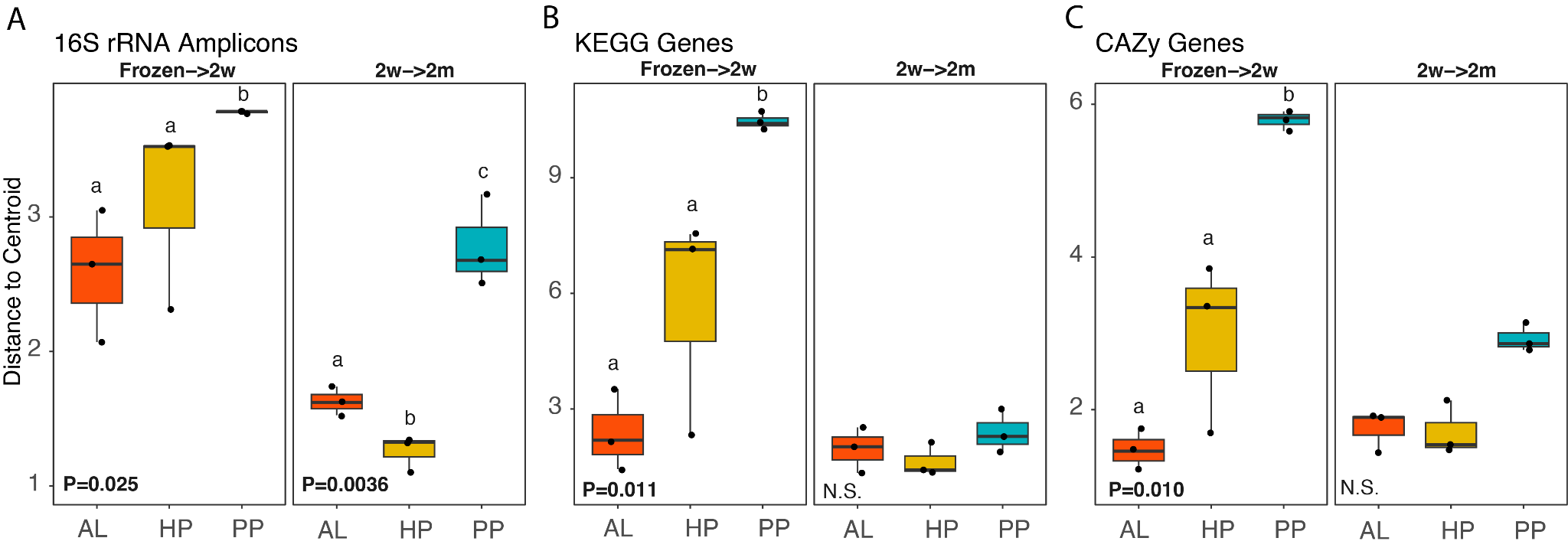

**Supplemental Figure 4. Magnitude and trajectory of changes in community and genetic structure for T_0_→two weeks and two weeks→two months.** For T_0_ to two weeks, data are the distances of thawed samples to the centroid of the frozen samples. For two weeks to two months, data are the distances of samples thawed for two months to the centroid of two week samples. Thawed samples represent cis inoculations (i.e., the inoculum and soil are the same). Differences among soils were assessed using a Kruskal-Wallis test followed by a Conover-Iman post hoc test.

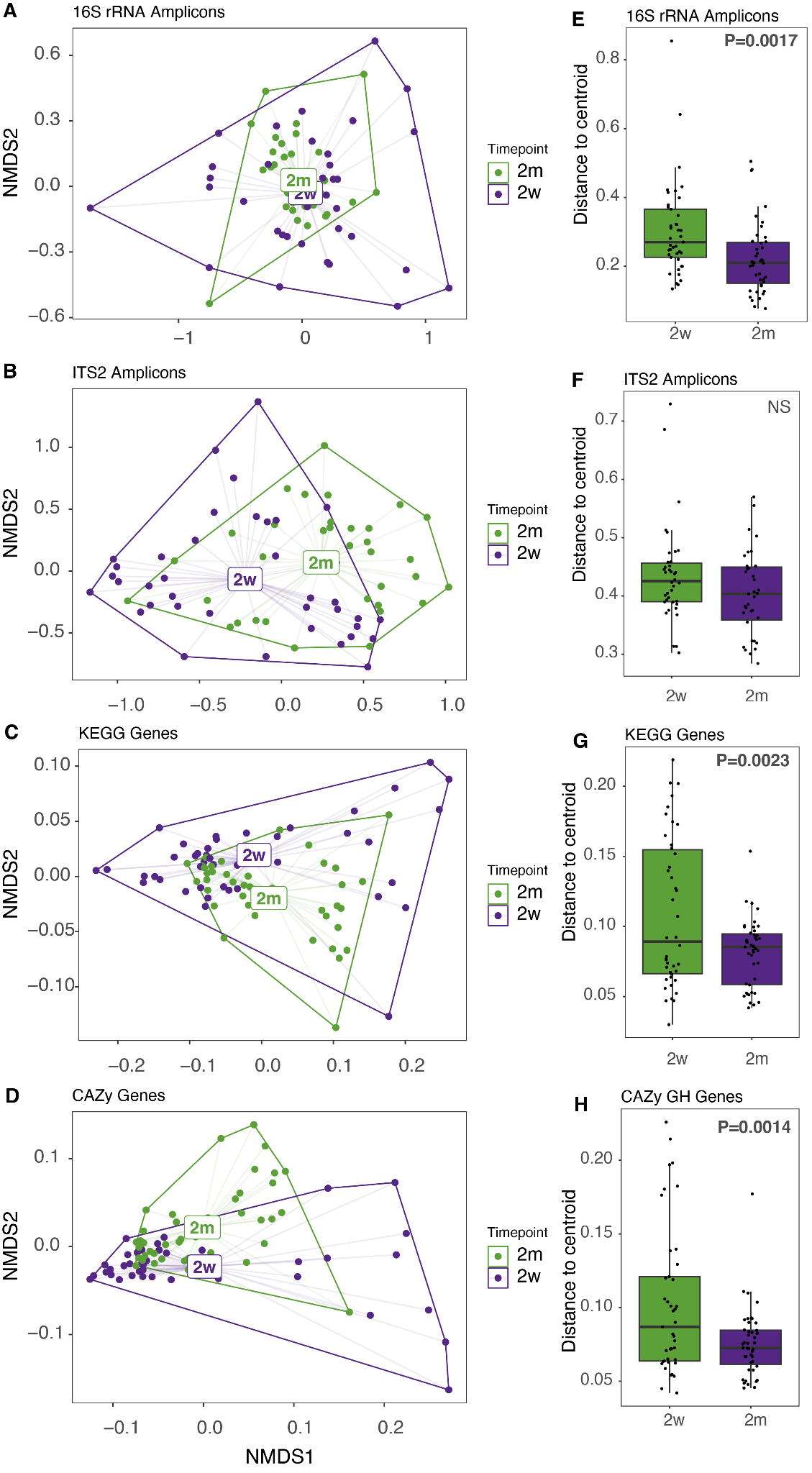

**Supplemental Figure 5. A-D)** Ordinations showing distance to centroid for each time point in amplicon and metagenome data. Data are visualized using NMDS plots based on Bray-Curtis dissimilarities. **E-H)** Differences in the distance to centroids between time points was tested using permutational analysis of multivariate dispersion and p-values are indicated in each panel

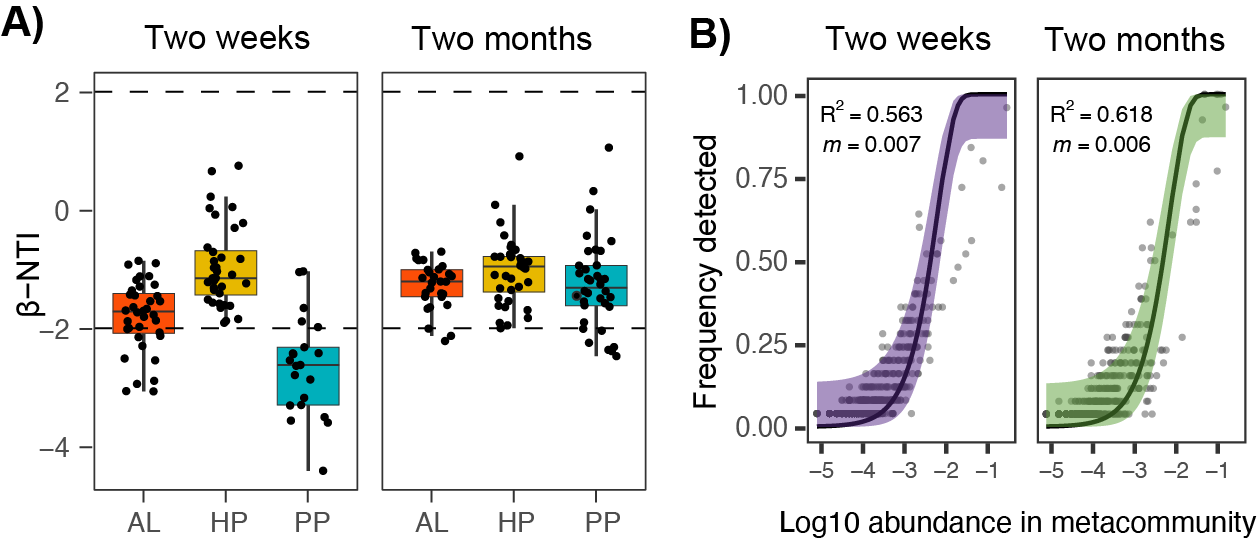

**Supplemental Figure 6. Phylogenetic turnover and abundance-occupancy dynamics during thaw progression. A)** β-nearest taxon index (βNTI) values in active layer (AL), Holocene permafrost (HP), and Pleistocene permafrost (PP) soils at two weeks and two months. **B)** Abundance-occupancy relationships showing the frequency of detection of taxa as a function of log₁₀ metacommunity abundance. Solid lines represent fits based on Sloan’s neutral community model, with corresponding R^2^ and migration parameter (m) values.

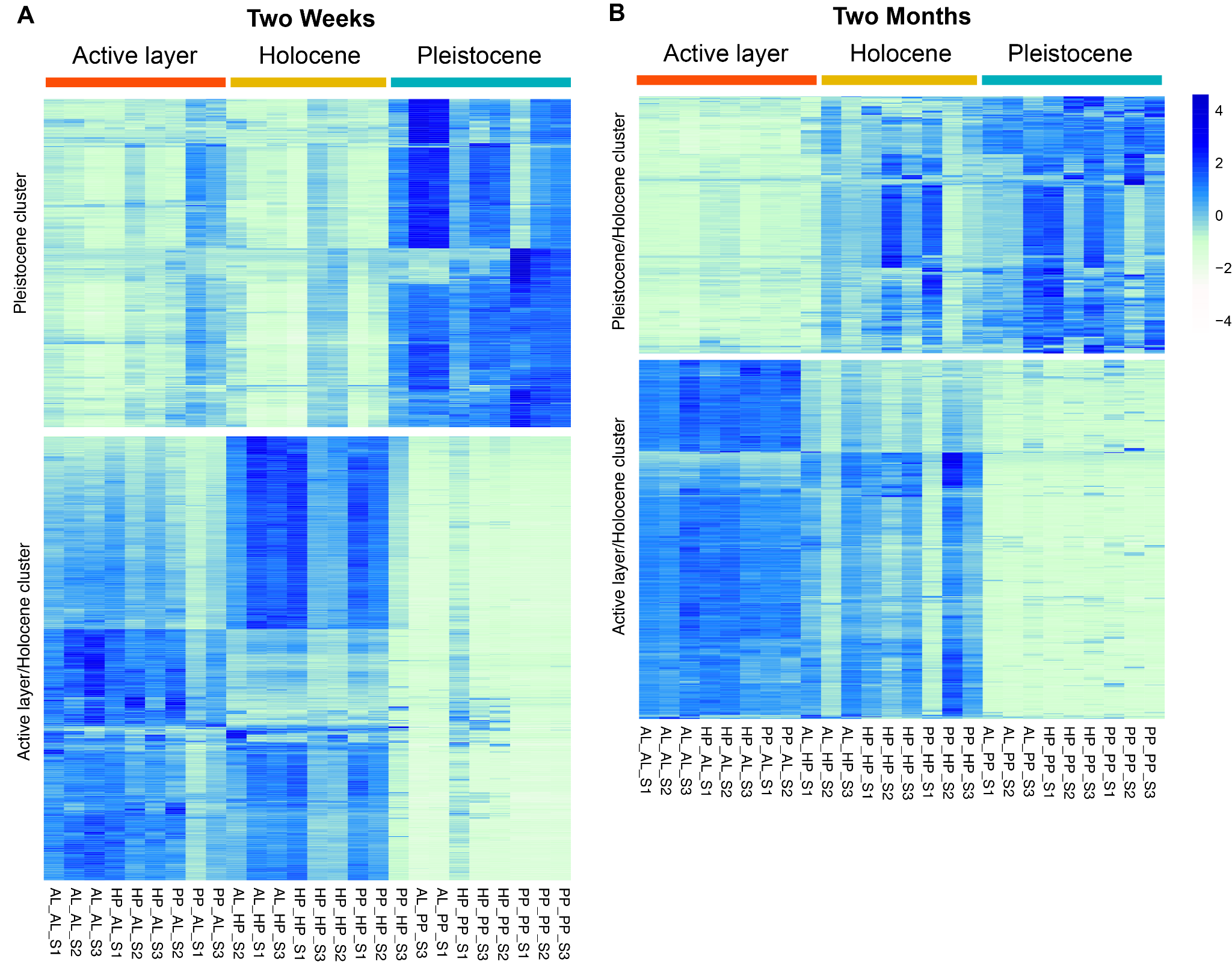

**Supplemental Figure 7. KEGG genes differing significantly among soil types.** Heatmaps show genes with significant differences in abundance among soils types after two weeks **(A)** and two months **(B)** of thaw. Genes are in rows and samples are in columns. Heatmap colors show the relative abundances of genes scaled by row. The division between genes more abundant in active layer soil/Holocene permafrost and Pleistocene permafrost is shown by large gaps in the heatmaps.

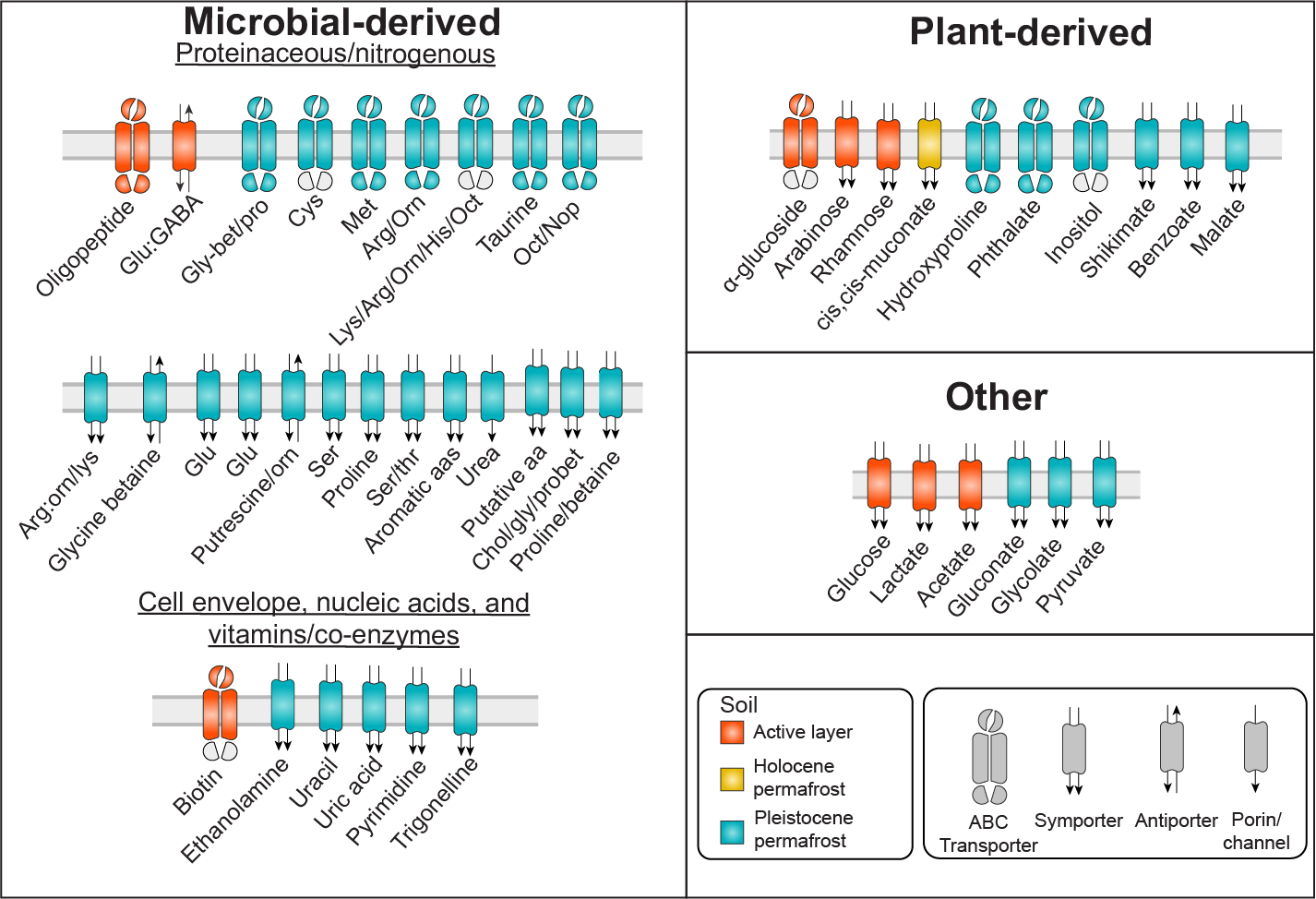

**Supplemental Figure 8. KEGG genes encoding carbon and nitrogen substrate transporters that are differentially abundant across soils after two months of thaw.** Placement into microbial-derived, plant-derived, and “other” categories is based on the likely dominant substrate source during early permafrost thaw. The microbial derived category is further divided into subcategories for proteinaceous or nitrogenous compounds, cell-envelope derived substrate, nucleic acids, and vitamins/co-enzymes. For transporters encoded by multiple genes, colored components represent the genes reaching significance. Amino acids are indicated by canonical three letter codes. Other abbreviations are in Fig 3. KEGG genes represented in the figure and corresponding descriptions are in Table S29.

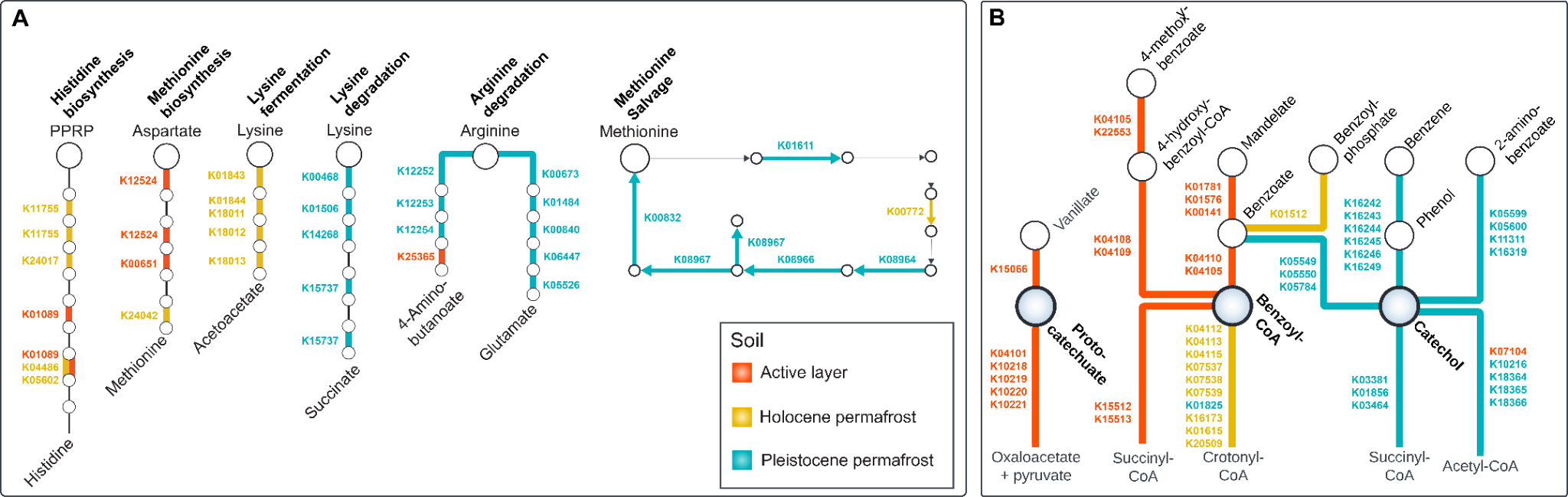

**Supplemental Figure 9.** KEGG pathway maps highlighting differentially abundant genes (DESeq2, *P* < 0.05). Color shows the soil with the highest abundance of each gene. KO numbers of differentially abundant genes are shown next to the corresponding pathway steps. **A)** Pathways related to amino acid metabolism. Bold colored lines show steps in the pathway represented by at least one differentially abundant gene. **B)** Pathways related to the degradation of aromatic compounds.

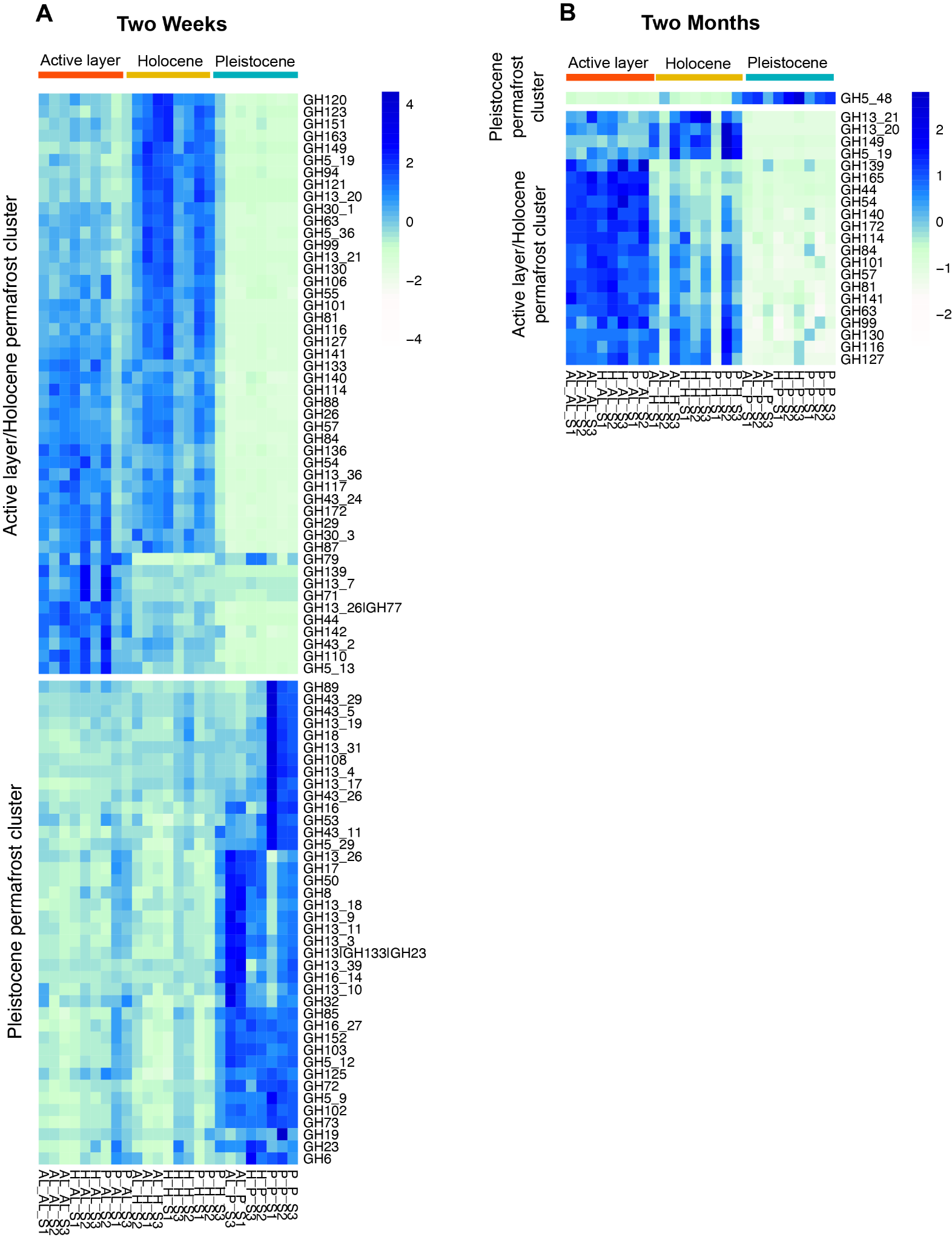

**Supplemental Figure 10. CAZy GH genes differing significantly among soil types.** Heatmaps show genes with significant differences in abundance (DESeq2, *P* < 0.05) among soils types after two weeks (A) and two months (B) of thaw. Genes are in rows and samples are in columns. Labels above the heatmap indicate soil. Sample names indicate the inoculum followed by the soil. Heatmap colors show the relative abundances of genes scaled by row. Clusters dividing active layer soil/Holocene permafrost and Pleistocene permafrost are shown by large gaps in the heatmaps.

### Supplemental Tables

**Supplemental Table 1.** Total carbon (%), total nitrogen (%), and C:N ratio for each soil type (active layer [AL], Holocene permafrost [HP], and Pleistocene permafrost [PP]) across timepoints (T_0_, 2 weeks [2w], and 2 months [2m]). Values are averages across replicates ± standard error.

| **Time** | **Soil** | **Inoculum** | **Total Carbon (%)** | **Total Nitrogen (%)** | **C:N** |
| --- | --- | --- | --- | --- | --- |
| T_0_ | AL |  | 3.91 ± 0.24 | 0.17 ± 0.02 | 23.17 ± 0.85 |
|  | HP |  | 2.98 ± 0.66 | 0.15 ± 0.02 | 19.95 ± 1.64 |
|  | PP |  | 3.92 ± 0.03 | 0.29 ± 0.003 | 13.48 ± 0.04 |
| 2w | AL | AL | 4.08 ± 0.22 | 0.23 ± 0.01 | 18.30 ± 1.73 |
|  |  | HP | 4.39 ± 0.50 | 0.20 ± 0.02 | 21.92 ± 0.35 |
|  |  | PP | 5.42 ± 1.19 | 0.23 ± 0.04 | 23.25 ± 1.21 |
|  | HP | AL | 2.21 ± 0.11 | 0.11 ± 0.01 | 20.23 ± 0.16 |
|  |  | HP | 2.23 ± 0.36 | 0.11 ± 0.004 | 20.46 ± 2.60 |
|  |  | PP | 1.96 ± 0.04 | 0.10 ± 0.003 | 19.06 ± 1.00 |
|  | PP | AL | 4.22 ± 0.10 | 0.28 ± 0.01 | 14.83 ± 0.07 |
|  |  | HP | 4.22 ± 0.21 | 0.27 ± 0.01 | 15.50 ± 1.03 |
|  |  | PP | 4.06 ± 0.13 | 0.28 ± 0.01 | 15.25 ± 0.21 |
| 2m | AL | AL | 3.89 ± 0.21 | 0.19 ± 0.01 | 20.73 ± 0.70 |
|  |  | HP | 3.75 ± 0.15 | 0.18 ± 0.01 | 22.47 ± 0.88 |
|  |  | PP | 6.33 ± 1.83 | 0.29 ± 0.11 | 22.55 ± 1.51 |
|  | HP | AL | 2.43 ± 0.21 | 0.12 ± 0.0006 | 20.96 ± 1.70 |
|  |  | HP | 3.98 ± 1.80 | 0.22 ± 0.10 | 18.02 ± 0.23 |
|  |  | PP | 2.52 ± 0.11 | 0.13 ± 0.01 | 19.78 ± 0.21 |
|  | PP | AL | 4.14 ± 0.08 | 0.27 ± 0.01 | 15.41 ± 0.59 |
|  |  | HP | 4.17 ± 0.15 | 0.27 ± 0.004 | 15.52 ± 0.41 |
|  |  | PP | 3.89 ± 0.05 | 0.27 ± 0.004 | 14.35 ± 0.14 |

**Supplemental Table 2.** Differences in total carbon, total nitrogen, and C:N among soil types at T_0_. One-way ANOVA results (F, p-values) and effect sizes (η²) are shown with Tukey-adjusted pairwise comparisons and Hedges’ g (95% confidence intervals). Soil types are active layer (AL), Holocene permafrost (HP), and Pleistocene permafrost (PP). The interpretation categories are based on Cohen (2013). ^136^.

|  | **ANOVA F (df=2,6)** | **ANOVA p-value** | **η²** | **Pairwise Comparison** | **Mean Diff** | **Tukey p** | **Hedges’ g (95% CI)** | **Interpretation** |
| --- | --- | --- | --- | --- | --- | --- | --- | --- |
| Total C (%) | 1.74 | 0.254 | 0.37 | HP – AL | -0.93 | 0.315 | — | ns |
|  |  |  |  | PP – AL | 0.01 | 0.999 | — | ns |
|  |  |  |  | PP – HP | 0.94 | 0.307 | — | ns |
| Total N (%) | 29.07 | **0.00082** | 0.91 | HP – AL | -0.023 | 0.524 | — | ns |
|  |  |  |  | PP – AL | 0.121 | **0.0024** | -5.23 [-8.92, -1.54] | Extreme |
|  |  |  |  | PP – HP | 0.145 | **0.0010** | -4.71 [-8.07, -1.33] | Extreme |
| C:N | 21.37 | **0.00186** | 0.88 | HP – AL | -3.23 | 0.162 | — | ns |
|  |  |  |  | PP – AL | -9.69 | **0.0016** | -7.36 [-12.43, -2.36] | Extreme |
|  |  |  |  | PP – HP | -6.46 | **0.0123** | -2.57 [-4.66,-0.39] | Very large |

**Supplemental Table 3**. Dissolved Organic Carbon (DOC) concentrations (mg/L) measured in leachates from active layer (AL), Holocene permafrost (HP), and Pleistocene permafrost (PP) at T0 (frozen), 2 weeks (2w), and 2 months (2m) following thaw. Replicate samples were pooled prior to leaching.

| **Sample ID** | **Soil** | **Time** | **DOC (mg/L)** |
| --- | --- | --- | --- |
| AL1_T0 | AL | T0 | 17.1 |
| AL3_T0 | AL | T0 | 15.5 |
| HP1_T0 | HP | T0 | 16.1 |
| HP2_T0 | HP | T0 | 16.1 |
| PP1_T0 | PP | T0 | 173.6 |
| PP2_T0 | PP | T0 | 172.6 |
| AL_PP_AL_2w | AL | 2w | 9.7 |
| AL_HP_AL_2w | AL | 2w | 11.8 |
| PP_AL_2w | AL | 2w | 23.9 |
| HP_AL_2w | AL | 2w | 14.8 |
| AL_AL_2w | AL | 2w | 13.9 |
| HP_HP_2w | HP | 2w | 13.08 |
| AL_HP_2w | HP | 2w | 10.5 |
| AL_HP_HP_2w | HP | 2w | 13.0 |
| PP_HP_2w | HP | 2w | 14.9 |
| HP_PP_HP_2w | HP | 2w | 22.8 |
| HP_PP_2w | PP | 2w | 42.4 |
| AL_PP_P_2w | PP | 2w | 54.8 |
| AL_PP_2w | PP | 2w | 44.5 |
| PP_PP_2w | PP | 2w | 42.8 |
| HP_PP_P_2w | PP | 2w | 36.5 |
| AL_AL_2m | AL | 2m | 4.8 |
| AL_HP_AL_2m | AL | 2m | 9.4 |
| PP_AL_2m | AL | 2m | 8.0 |
| AL_PP_AL_2m | AL | 2m | 8.2 |
| HP_AL_2m | AL | 2m | 4.8 |
| PP_HP_2m | HP | 2m | 17.6 |
| HP_HP_2m | HP | 2m | 11.5 |
| AL_HP_HP_2m | HP | 2m | 13.6 |
| HP_PP_HP_2m | HP | 2m | 14.5 |
| AL_HP_2m | HP | 2m | 9.2 |
| AL_PP_2m | PP | 2m | 37.9 |
| HP_PP_P_2m | PP | 2m | 37.4 |
| AL_PP_P_2m | PP | 2m | 42.8 |
| PP_PP_2m | PP | 2m | 20.8 |

**Supplemental Table 4**. Low molecular weight organic acids (mM) measured in leachates in T_0_ samples from active layer (AL), Holocene permafrost (HP), and Pleistocene permafrost (PP) and thawed Pleistocene permafrost samples.

| **Sample ID** | **Acetate** | **Butyrate** | **Propionate** | **Timepoint** | **Inoculum** | **Soil** |
| --- | --- | --- | --- | --- | --- | --- |
| AL1 | 0 | 0.555 | 0 | T0 |  | AL |
| AL2 | 0 | 0.39 | 0 | T0 |  | AL |
| AL3 | 0 | 0.345 | 0 | T0 |  | AL |
| HP1 | 0 | 0 | 0 | T0 |  | HP |
| HP2 | 0 | 0 | 0 | T0 |  | HP |
| HP3 | 0.15 | 0 | 0 | T0 |  | HP |
| PP1 | 2.43 | 15.78 | 2.565 | T0 |  | PP |
| PP2 | 2.58 | 14.775 | 2.595 | T0 |  | PP |
| PP3 | 4.455 | 29.085 | 3.69 | T0 |  | PP |
| AL_PP_S2_2w | 0.27 | 1.215 | 0.855 | 2w | AL | PP |
| AL_PP_S3_2w | 0 | 0.87 | 0.465 | 2w | AL | PP |
| HP_PP_S1_2w | 0.36 | 0.36 | 0.855 | 2w | HP | PP |
| HP_PP_S2_2w | 0.225 | 0.54 | 0.87 | 2w | HP | PP |
| HP_PP_S3_2w | 0.165 | 0.87 | 4.14 | 2w | HP | PP |
| PP_PP_S1_2w | 0.195 | 0.6 | 0.87 | 2w | PP | PP |
| PP_PP_S2_2w | 0.165 | 0.81 | 0 | 2w | PP | PP |
| PP_PP_S3_2w | 0.255 | 0.405 | 0.855 | 2w | PP | PP |
| AL_PP_S1_2m | 0.315 | 0.225 | 0.855 | 2m | AL | PP |
| AL_PP_S2_2m | 0.525 | 0 | 0.855 | 2m | AL | PP |
| AL_PP_S3_2m | 0.15 | 0 | 0.87 | 2m | AL | PP |
| HP_PP_S1_2m | 0.21 | 0 | 0.87 | 2m | HP | PP |
| HP_PP_S2_2m | 0.24 | 0 | 0 | 2m | HP | PP |
| PP_PP_S1_2m | 0.315 | 0 | 0 | 2m | PP | PP |
| PP_PP_S2_2m | 0.165 | 0 | 0 | 2m | PP | PP |
| PP_PP_S3_2m | 0.21 | 0 | 0 | 2m | PP | PP |

**Supplemental Table 5. FT-ICR-MS formulae**

[Table S5](https://docs.google.com/spreadsheets/d/15vlx8NTy51j6IS1eyxyVOu-rg54ghOdejO7mqJXD8hE/edit?usp=sharing)

**Supplementary Table 6. FT-ICR-MS summary metrics**

[Table S6.xlsx](https://docs.google.com/spreadsheets/d/1YwfVBHqorYrEWLvoC1yFXxpVcQCDIGtc/edit?usp=sharing&ouid=102681131962325051838&rtpof=true&sd=true)

**Supplemental Tables 7-10. Nodes from quadripartite networks**

[SupplementaryTables7-10.xlsx](https://docs.google.com/spreadsheets/d/1ZSSYG2pdVymyPX8vxD-wP5BBugAd_lsI/edit?usp=drive_link&ouid=102681131962325051838&rtpof=true&sd=true)

**Supplemental Table 11. PERMANOVA, PERMDISP, and dbRDA results**

| **PERMANOVA** | | | | | |
| --- | --- | --- | --- | --- | --- |
|  | df | SS | pseudoF | R^2^ | P-value |
| **16S rRNA Amplicons** |  |  |  |  |  |
| Inoculum | 2 | 0.8545 | 4.8398 | 0.08865 | **0.0001** |
| Time point | 1 | 1.4496 | 16.4210 | 0.15038 | **0.0001** |
| Soil | 2 | 2.4980 | 14.1486 | 0.25915 | **0.0001** |
| Inoculum x time point | 2 | 0.3866 | 2.1899 | 0.04011 | **0.0065** |
| Inoculum x soil origin | 4 | 0.5315 | 1.5051 | 0.05513 | **0.0463** |
| Time point x soil origin | 2 | 0.5858 | 3.3178 | 0.06077 | **0.0001** |
| Inoculum x time point x soil origin | 4 | 0.4202 | 1.1901 | 0.04360 | 0.2259 |
| **ITS2 Amplicons** |  |  |  |  |  |
| Inoculum | 2 | 1.1627 | 4.3201 | 0.14325 | **0.0002** |
| Time point | 1 | 0.5686 | 4.2256 | 0.07006 | **0.0004** |
| Soil | 2 | 0.9029 | 3.3549 | 0.11125 | **0.0002** |
| Inoculum x time point | 2 | 0.2531 | 0.9336 | 0.03096 | 0.5215 |
| Inoculum x soil origin | 4 | 0.6316 | 1.1734 | 0.07782 | 0.2515 |
| Time point x soil origin | 2 | 0.8726 | 3.2422 | 0.10751 | **0.0002** |
| Inoculum x time point x soil origin | 4 | 0.3624 | 0.8976 | 0.04465 | 0.6043 |
| **KEGG Genes** |  |  |  |  |  |
| Inoculum | 2 | 0.02360 | 3.1107 | 0.03803 | **0.0232** |
| Time point | 1 | 0.04327 | 11.4075 | 0.06973 | **0.0002** |
| Soil origin | 2 | 0.32095 | 42.3048 | 0.51721 | **0.0001** |
| Inoculum x time point | 2 | 0.01522 | 2.0062 | 0.02453 | 0.1008 |
| Inoculum x soil origin | 4 | 0.02321 | 1.5297 | 0.03740 | 0.1592 |
| Time point x soil origin | 2 | 0.04892 | 6.4479 | 0.07883 | **0.0004** |
| Inoculum x time point x soil origin | 4 | 0.01640 | 1.0809 | 0.02643 | 0.3785 |
| **CAZy Genes** |  |  |  |  |  |
| Inoculum | 2 | 0.01966 | 3.1204 | 0.03693 | **0.0177** |
| Time point | 1 | 0.04425 | 14.0511 | 0.08341 | **0.0001** |
| Soil origin | 2 | 0.23846 | 37.8556 | 0.44797 | **0.0001** |
| Inoculum x time point | 2 | 0.01929 | 3.0623 | 0.03624 | **0.0180** |
| Inoculum x soil origin | 4 | 0.02147 | 1.7041 | 0.04033 | 0.0935 |
| Time point x soil origin | 2 | 0.06038 | 9.5851 | 0.11343 | **0.0001** |
| Inoculum x time point x soil origin | 4 | 0.10708 | 1.7242 | 0.04081 | 0.0870 |
| **PERMDISP** | | | | | |
|  | df | SS | MeanSq | F | P-value |
| **KEGG Genes** |  |  |  |  |  |
| Soil | 2 | 0.01273 | 0.00637 | 6.46174 | **0.0026** |
| Time point | 1 | 0.00904 | 0.00904 | 4.37688 | **0.0401** |
| Inoculum | 2 | 0.00143 | 0.00072 | 0.30595 | 0.7438 |
| **CAZy Genes** |  |  |  |  |  |
| Soil | 2 | 0.02764 | 0.01381 | 19.50311 | **0.0001** |
| Time point | 1 | 0.00839 | 0.00839 | 3.46761 | 0.0690 |
| Inoculum | 2 | 0.00432 | 0.00216 | 0.86879 | 0.0690 |
| **dbRDA** | | | | | |
|  |  | df | SS | F | P-value |
| **KEGG Genes** | Soil | 2 | 0.32095 | 31.72 | **0.0001** |
| **CAZy Genes** | Soil | 2 | 0.23864 | 28.852 | **0.0001** |

df: degrees of freedom, SS: sum of squares, Pseudo-F: F value by permutation, P: *P-*values based on 9999 permutations calculated on a Bray-Curtis dissimilarity matrix. Bold numbers are *P*-values < 0.05.

**Supplemental Table 12. PERMANOVA results for two-week and two-month time points.**

| **Two Weeks** | | | | | |
| --- | --- | --- | --- | --- | --- |
|  | df | SS | pseudoF | R^2^ | Pr(>F) |
| **16S rRNA Amplicons** |  |  |  |  |  |
| Inoculum | 2 | 0.8268 | 4.1652 | 0.15832 | **0.0004** |
| Soil | 2 | 2.1469 | 10.8150 | 0.41107 | **0.0001** |
| Inoculum x Soil | 4 | 0.6609 | 1.6646 | 0.12654 | **0.0499** |
| **ITS2 Amplicons** |  |  |  |  |  |
| Inoculum | 2 | 0.5918 | 1.9292 | 0.12129 | **0.0347** |
| Soil | 2 | 1.3872 | 4.5217 | 0.28428 | **0.0001** |
| Inoculum x Soil | 4 | 0.7532 | 1.2275 | 0.15435 | 0.2095 |
| **KEGG Genes** |  |  |  |  |  |
| Inoculum | 2 | 0.02866 | 3.1255 | 0.07343 | **0.0472** |
| Soil | 2 | 0.25016 | 27.2760 | 0.64087 | **0.0001** |
| Inoculum x Soil | 4 | 0.03356 | 1.8298 | 0.08589 | 0.1338 |
| **CAZy Genes** |  |  |  |  |  |
| Inoculum | 2 | 0.02485 | 4.0269 | 0.07487 | **0.0186** |
| Soil | 2 | 0.22063 | 35.7562 | 0.66482 | **0.0001** |
| Inoculum x Soil | 4 | 0.03394 | 2.7501 | 0.10227 | **0.0315** |
| **Two Months** | | | | | |
|  | df | SS | pseudoF | R^2^ | Pr(>F) |
| **16S rRNA Amplicons** |  |  |  |  |  |
| Inoculum | 2 | 0.4703 | 2.4707 | 0.13212 | **0.0010** |
| Soil | 2 | 1.0828 | 5.6881 | 0.30416 | **0.0001** |
| Inoculum x Soil | 4 | 0.3887 | 1.0210 | 0.10919 | 0.4363 |
| **ITS2 Amplicons** |  |  |  |  |  |
| Inoculum | 2 | 0.79146 | 3.1963 | 0.27623 | **0.0043** |
| Soil | 2 | 0.45495 | 1.8373 | 0.15879 | 0.0719 |
| Inoculum x Soil | 4 | 0.25698 | 0.6919 | 0.08966 | 0.8111 |
| **KEGG Genes** |  |  |  |  |  |
| Inoculum | 2 | 0.011198 | 1.8765 | 0.05918 | 0.1186 |
| Soil | 2 | 0.120065 | 20.1194 | 0.63447 | **0.0001** |
| Inoculum x Soil | 4 | 0.007248 | 0.6073 | 0.03830 | 0.7859 |
| **CAZy Genes** |  |  |  |  |  |
| Inoculum | 2 | 0.012407 | 2.0051 | 0.08214 | 0.0690 |
| Soil | 2 | 0.076307 | 12.3322 | 0.50520 | **0.0001** |
| Inoculum x Soil | 4 | 0.009736 | 0.7867 | 0.06446 | 0.6698 |

df: degrees of freedom, SS: sum of squares, Pseudo-F: F value by permutation, Pr(>F): *P-*values based on 9999 permutations calculated on a Bray-Curtis dissimilarity matrix. Bold numbers are *P*-values < 0.05.

**Supplemental Table 13. Permutational analysis of multivariate dispersion (PERMDISP) across timepoints**

|  | **df** | **SS** | **Mean Sq** | **F** | **Pr(>F)** |
| --- | --- | --- | --- | --- | --- |
| 16S rRNA amplicons | 1 | 0.1396 | 0.13904 | 9.6635 | **0.0023** |
| ITS2 amplicons | 1 | 0.01841 | 0.0184121 | 2.9695 | 0.0931 |
| KEGG genes | 1 | 0.016373 | 0.016373 | 9.6713 | **0.0023** |
| CAZy GH genes | 1 | 0.015265 | 0.0152646 | 10.155 | **0.0014** |

df: degrees of freedom, SS: sum of squares, Mean Sq: mean square. P values were calculated using 9999 permutations. Bold numbers are *P*-values < 0.05

**Supplemental Table 14. Differentially abundant ASVs after two weeks and two months of thaw**

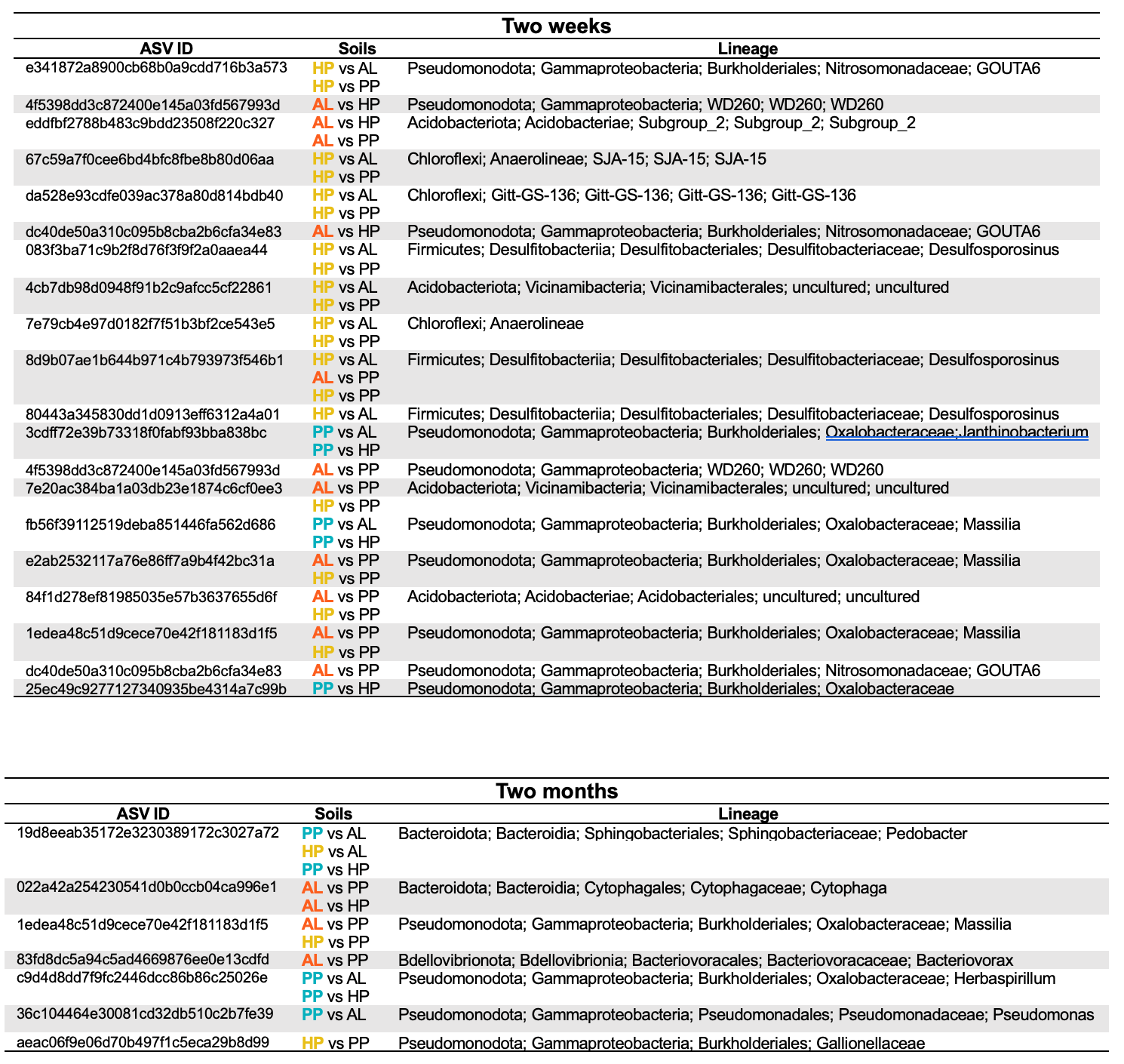

Differentially abundant amplicon sequence variants (ASVs) were identified using DESeq2 (*P* < 0.05). The soils column specifies significant comparisons, with the first soil (bolded and colored) indicating the soil with the higher abundance. AL (red): active layer, HP (yellow): Holocene permafrost, PP (blue): Pleistocene permafrost. ASV lineages are phylum, class, order, family, genus.

**Supplemental Tables 15-17. DESeq2 results for 16S rRNA ASVs at two weeks post thaw.**

[**SupplementalTables15-17.xlsx**](https://docs.google.com/spreadsheets/d/1jX56k1w8KowElNXc66PsmnXdBTPT0GS_/edit?usp=sharing&ouid=102681131962325051838&rtpof=true&sd=true)

**Supplemental Tables 18-20. DESeq2 results for 16S rRNA ASVs at two months post thaw.**

[**SupplementalTables18-20.xlsx**](https://docs.google.com/spreadsheets/d/1MQKPWpjnxjPSxqe8OvgO6ORye6dsKf2K/edit?usp=sharing&ouid=102681131962325051838&rtpof=true&sd=true)

**Supplemental Tables 21-23. DESeq2 results for KEGG genes at two weeks post thaw.**

[**SupplementalTables21-23.xlsx**](https://docs.google.com/spreadsheets/d/1KQaslQdA73vPQ31y8rCOpLvyQzunlhk3/edit?usp=sharing&ouid=102681131962325051838&rtpof=true&sd=true)

**Supplemental Tables 24-26. DESeq2 results for KEGG genes at two months post thaw.**

[SupplementalTables24-26.xlsx](https://docs.google.com/spreadsheets/d/1baahFqFDiYelthAPfMXx-HysKC3a3XF-/edit?usp=sharing&ouid=102681131962325051838&rtpof=true&sd=true)

**Supplemental Tables 27-28. KEGG transport genes related to the uptake of carbon and nitrogen substrate that are differentially abundant across soils**

[**Supplemental Tables 27-28.xlsx**](https://docs.google.com/spreadsheets/d/10eejnm8dxARjTarCC0zrhjwpuSDhW18O/edit?usp=sharing&ouid=102681131962325051838&rtpof=true&sd=true)

**Supplemental Tables 29-31. DESeq2 results for CAZy genes at two weeks post thaw.**

[**SupplementalTables29-31.xlsx**](https://docs.google.com/spreadsheets/d/1npBRxl0k7qYLGhkU__NkXZhUpxdcsTrH/edit?usp=sharing&ouid=102681131962325051838&rtpof=true&sd=true)

**Supplemental Tables 32-34. DESeq2 results for CAZy genes at two months post thaw.**

[SupplementalTables32-34.xlsx](https://docs.google.com/spreadsheets/d/1mq7VUKa4W1IUkkfuP8PYWw8iX_ajtv0d/edit?usp=sharing&ouid=102681131962325051838&rtpof=true&sd=true)

### References

### 1. Zimov, S. A., Schuur, E. A. & Chapin, F. S. Climate change. Permafrost and the global carbon budget. *Science (New York, NY)* vol. 312 1612–1613 Preprint at https://doi.org/10.1126/science.1128908 (2006).

### 2. Chadburn, S. E. *et al.* An observation-based constraint on permafrost loss as a function of global warming. *Nat. Clim. Chang.* 7, 340–344 (2017).

### 3. Tarnocai, C. *et al.* Soil organic carbon pools in the northern circumpolar permafrost region: SOIL ORGANIC CARBON POOLS. *Global Biogeochem. Cycles* 23, (2009).

### 4. Schuur, E. A. G. *et al.* Climate change and the permafrost carbon feedback. *Nature* 520, 171–179 (2015).

### 5. Zimov, N. S. *et al.* Carbon storage in permafrost and soils of the mammoth tundra-steppe biome: Role in the global carbon budget. *Geophys. Res. Lett.* 36, (2009).

### 6. Zimov, S. A., Zimov, N. S., Tikhonov, A. N. & Chapin, F. S., III. Mammoth steppe: a high-productivity phenomenon. *Quat. Sci. Rev.* 57, 26–45 (2012).

### 7. Schirrmeister, L., Froese, D., Tumskoy, V., Grosse, G. & Wetterich, S. PERMAFROST AND PERIGLACIAL FEATURES | yedoma: Late Pleistocene ice-rich syngenetic permafrost of Beringia. in *Encyclopedia of Quaternary Science* 542–552 (Elsevier, 2013).

### 8. Kuhry, P. *et al.* Characterisation of the permafrost carbon pool: Permafrost carbon. *Permafrost Periglacial Processes* 24, 146–155 (2013).

### 9. Wetterich, S. *et al.* Ice Complex formation in arctic East Siberia during the MIS3 Interstadial. *Quaternary Science Reviews* 84, 39–55 (2014).

### 10. Martens, J. *et al.* Stabilization of mineral-associated organic carbon in Pleistocene permafrost. *Nat. Commun.* 14, 2120 (2023).

### 11. Strauss, J. *et al.* Organic-matter quality of deep permafrost carbon – a study from Arctic Siberia. *Biogeosciences* 12, 2227–2245 (2015).

### 12. Schirrmeister, L. *et al.* Fossil organic matter characteristics in permafrost deposits of the northeast Siberian Arctic. *J. Geophys. Res.* 116, (2011).

### 13. Drake, T. W., Wickland, K. P., Spencer, R. G. M., McKnight, D. M. & Striegl, R. G. Ancient low-molecular-weight organic acids in permafrost fuel rapid carbon dioxide production upon thaw. *Proc. Natl. Acad. Sci. U. S. A.* 112, 13946–13951 (2015).

### 14. Vonk, J. E. *et al.* High biolability of ancient permafrost carbon upon thaw: BIOLABILITY OF ANCIENT PERMAFROST CARBON. *Geophys. Res. Lett.* 40, 2689–2693 (2013).

### 15. Strauss, J. *et al.* Deep Yedoma permafrost: A synthesis of depositional characteristics and carbon vulnerability. *Earth-Sci. Rev.* 172, 75–86 (2017).

### 16. Holm, S. *et al.* Methanogenic response to long-term permafrost thaw is determined by paleoenvironment. *FEMS Microbiol. Ecol.* 96, fiaa021 (2020).

### 17. Oswald, W. W., Brubaker, L. B., Hu, F. S. & Kling, G. W. Holocene pollen records from the central Arctic Foothills, northern Alaska: testing the role of substrate in the response of tundra to climate change. *J. Ecol.* 91, 1034–1048 (2003).

### 18. Mann, D. H., Peteet, D. M., Reanier, R. E. & Kunz, M. L. Responses of an arctic landscape to Lateglacial and early Holocene climatic changes: the importance of moisture. *Quat. Sci. Rev.* 21, 997–1021 (2002).

### 19. Ernakovich, J. G. *et al.* Microbiome assembly in thawing permafrost and its feedbacks to climate. *Glob. Chang. Biol.* 28, 5007–5026 (2022).

### 20. Waldrop, M. P. *et al.* Permafrost microbial communities and functional genes are structured by latitudinal and soil geochemical gradients. *ISME J.* 17, 1224–1235 (2023).

### 21. Mackelprang, R. *et al.* Microbial survival strategies in ancient permafrost: insights from metagenomics. *ISME J.* 11, 2305–2318 (2017).

### 22. Waldrop, M. P. *et al.* Microbial ecology of permafrost soils: Populations, processes, and perspectives. *Permafrost Periglacial Processes* 36, 245–258 (2025).

### 23. Monteux, S. *et al.* Long-term in situ permafrost thaw effects on bacterial communities and potential aerobic respiration. *ISME J.* 12, 2129–2141 (2018).

### 24. Martiny, J. B. *et al.* Microbial legacies alter decomposition in response to simulated global change. *ISME J.* 11, 490–499 (2017).

### 25. Albright, M. B. N. & Martiny, J. B. H. Dispersal alters bacterial diversity and composition in a natural community. *ISME J.* 12, 296–299 (2018).

### 26. Spencer, R. G. M. *et al.* Detecting the signature of permafrost thaw in Arctic rivers. *Geophys. Res. Lett.* 42, 2830–2835 (2015).

### 27. D’Andrilli, J. *et al.* Advancing chemical lability assessments of organic matter using a synthesis of FT-ICR MS data across diverse environments and experiments. *Org. Geochem.* 184, 104667 (2023).

### 28. Textor, S. R., Wickland, K. P., Podgorski, D. C., Johnston, S. E. & Spencer, R. G. M. Dissolved Organic Carbon Turnover in Permafrost-Influenced Watersheds of Interior Alaska: Molecular Insights and the Priming Effect. *Front Earth Sci. Chin.* 7, 275 (2019).

### 29. Belova, S. E. *et al.* Hydrolytic capabilities as a key to environmental success: Chitinolytic and cellulolytic Acidobacteria from acidic sub-arctic soils and boreal peatlands. *Front. Microbiol.* 9, 2775 (2018).

### 30. Kalam, S. *et al.* Recent understanding of soil Acidobacteria and their ecological significance: A critical review. *Front. Microbiol.* 11, 580024 (2020).

### 31. Yang, Y., Chen, Q., Zhou, Y., Yu, W. & Shi, Z. Soil bacterial community composition and function play roles in soil carbon balance in alpine timberline ecosystems. *J. Soils Sediments* 24, 323–336 (2024).

### 32. Ivanova, A. A., Zhelezova, A. D., Chernov, T. I. & Dedysh, S. N. Linking ecology and systematics of acidobacteria: Distinct habitat preferences of the Acidobacteriia and Blastocatellia in tundra soils. *PLoS One* 15, e0230157 (2020).

### 33. Sipes, K. *et al.* Permafrost active layer microbes from NY Ålesund, Svalbard (79°N) show autotrophic and heterotrophic metabolisms with diverse carbon-degrading enzymes. *Front. Microbiol.* 12, 757812 (2021).

### 34. Sipes, K. *et al.* Eight Metagenome-Assembled Genomes Provide Evidence for Microbial Adaptation in 20,000- to 1,000,000-Year-Old Siberian Permafrost. *Appl. Environ. Microbiol.* 87, e0097221 (2021).

### 35. O’Brien, J. M. *et al.* Consistent microorganisms respond during aerobic thaw of Alaskan permafrost soils. *bioRxiv* (2025) doi:10.1101/2025.08.11.669168.

### 36. Rice, A. V. & Currah, R. S. Oidiodendron: A survey of the named species and related anamorphs of Myxotrichum. *Stud. Mycol.* 53, 83–120 (2005).

### 37. Sigler, L., Lumley, T. C. & Currah, R. S. New species and records of saprophytic ascomycetes (Myxotrichaceae) from decaying logs in the boreal forest. *Mycoscience* 41, 495–502 (2000).

### 38. Maillard, F., Schilling, J., Andrews, E., Schreiner, K. M. & Kennedy, P. Functional convergence in the decomposition of fungal necromass in soil and wood. *FEMS Microbiol. Ecol.* 96, (2020).

### 39. Leifheit, E. F. *et al.* Fungal traits help to understand the decomposition of simple and complex plant litter. *FEMS Microbiol. Ecol.* 100, (2024).

### 40. Li, F. *et al.* The changes of chemical molecular components in soil organic matter are associated with fungus Mortierella capitata K. *Soil Tillage Res.* 227, 105598 (2023).

### 41. Fukasawa, Y., Osono, T. & Takeda, H. Wood decomposing abilities of diverse lignicolous fungi on nondecayed and decayed beech wood. *Mycologia* 103, 474–482 (2011).

### 42. Gundersen, M. J. S. & Vadstein, O. An exploration of the impact of phylogenetic tree structure on NTI and βNTI estimates of community assembly. *Sci. Rep.* 14, 23480 (2024).

### 43. Bethencourt, L. *et al.* Genome reconstruction reveals distinct assemblages of Gallionellaceae in surface and subsurface redox transition zones. *FEMS Microbiol. Ecol.* 96, (2020).

### 44. Hoover, R. L., Keffer, J. L., Polson, S. W. & Chan, C. S. Gallionellaceae pangenomic analysis reveals insight into phylogeny, metabolic flexibility, and iron oxidation mechanisms. *mSystems* 8, e0003823 (2023).

### 45. Kowalchuk, G. A. & Stephen, J. R. Ammonia-oxidizing bacteria: a model for molecular microbial ecology. *Annu. Rev. Microbiol.* 55, 485–529 (2001).

### 46. Fierer, N., Bradford, M. A. & Jackson, R. B. Toward an ecological classification of soil bacteria. *Ecology* 88, 1354–1364 (2007).

### 47. Kumar, A., Männistö, M. K., Kerkhof, L. J. & Häggblom, M. M. Genome analysis reveals diversity and functional potential of novel Janthinobacterium species from subarctic soils. *Microbiologyopen* 15, e70239 (2026).

### 48. Straub, D., Rothballer, M., Hartmann, A. & Ludewig, U. The genome of the endophytic bacterium H. frisingense GSF30(T) identifies diverse strategies in the Herbaspirillum genus to interact with plants. *Front. Microbiol.* 4, 168 (2013).

### 49. Eilers, K. G., Lauber, C. L., Knight, R. & Fierer, N. Shifts in bacterial community structure associated with inputs of low molecular weight carbon compounds to soil. *Soil Biol. Biochem.* 42, 896–903 (2010).

### 50. Werdin-Pfisterer, N. R., Kielland, K. & Boone, R. D. Soil amino acid composition across a boreal forest successional sequence. *Soil Biol. Biochem.* 41, 1210–1220 (2009).

### 51. Andresen, L. C. *et al.* Patterns of free amino acids in tundra soils reflect mycorrhizal type, shrubification, and warming. *Mycorrhiza* 32, 305–313 (2022).

### 52. Akashi, H. & Gojobori, T. Metabolic efficiency and amino acid composition in the proteomes of Escherichia coli and Bacillus subtilis. *Proc. Natl. Acad. Sci. U. S. A.* 99, 3695–3700 (2002).

### 53. Wilkinson, A., Hill, P. W., Farrar, J. F., Jones, D. L. & Bardgett, R. D. Rapid microbial uptake and mineralization of amino acids and peptides along a grassland productivity gradient. *Soil Biol. Biochem.* 72, 75–83 (2014).

### 54. Ma, Q. *et al.* Substrate control of sulphur utilisation and microbial stoichiometry in soil: Results of 13C, 15N, 14C, and 35S quad labelling. *ISME J.* 15, 3148–3158 (2021).

### 55. Fuchs, G., Boll, M. & Heider, J. Microbial degradation of aromatic compounds - from one strategy to four. *Nat. Rev. Microbiol.* 9, 803–816 (2011).

### 56. Masai, E., Katayama, Y. & Fukuda, M. Genetic and biochemical investigations on bacterial catabolic pathways for lignin-derived aromatic compounds. *Biosci. Biotechnol. Biochem.* 71, 1–15 (2007).

### 57. Harwood, C. S. & Parales, R. E. The beta-ketoadipate pathway and the biology of self-identity. *Annu. Rev. Microbiol.* 50, 553–590 (1996).

### 58. Vonk, J. E. *et al.* Dissolved organic carbon loss from Yedoma permafrost amplified by ice wedge thaw. *Environmental Research Letters* vol. 8 035023 Preprint at https://doi.org/10.1088/1748-9326/8/3/035023 (2013).

### 59. Leewis, M.-C. *et al.* Life at the Frozen Limit: Microbial Carbon Metabolism Across a Late Pleistocene Permafrost Chronosequence. *Front. Microbiol.* 11, 1753 (2020).

### 60. Doherty, S. J., Thurston, A. K. & Barbato, R. A. Active layer and permafrost microbial community coalescence increases soil activity and diversity in mixed communities compared to permafrost alone. *Front. Microbiol.* 16, 1579156 (2025).

### 61. Geers, A. U., Buijs, Y., Schostag, M. D., Elberling, B. & Bentzon-Tilia, M. Exploring the biosynthesis potential of permafrost microbiomes. *Environ. Microbiome* 19, 96 (2024).

### 62. Varsadiya, M., Urich, T., Hugelius, G. & Bárta, J. Microbiome structure and functional potential in permafrost soils of the Western Canadian Arctic. *FEMS Microbiol. Ecol.* 97, fiab008 (2021).

### 63. Weedon, J. T. *et al.* Compositional stability of the bacterial community in a climate-sensitive sub-arctic peatland. *Front. Microbiol.* 8, 317 (2017).

### 64. O’Brien, J. M. *et al.* Consistent microbial responses during the aerobic thaw of Alaskan permafrost soils. *Front. Microbiol.* 16, 1654065 (2025).

### 65. Chen, S. *et al.* Divergent responses of bacterial communities to permafrost degradation and their associations with carbon across vertical profiles. *Adv. Sci. (Weinh.)* e10516 (2026).

### 66. Louca, S. *et al.* Function and functional redundancy in microbial systems. *Nat. Ecol. Evol.* 2, 936–943 (2018).

### 67. Soufi, H. H. *et al.* Taxonomic variability and functional stability across Oregon coastal subsurface microbiomes. *Commun. Biol.* 7, 1663 (2024).

### 68. Burke, C., Steinberg, P., Rusch, D., Kjelleberg, S. & Thomas, T. Bacterial community assembly based on functional genes rather than species. *Proc. Natl. Acad. Sci. U. S. A.* 108, 14288–14293 (2011).

### 69. Allison, S. D. & Martiny, J. B. H. Resistance, resilience, and redundancy in microbial communities. *Proceedings of the National Academy of Sciences* vol. 105 11512–11519 Preprint at https://doi.org/10.1073/pnas.0801925105 (2008).

### 70. Shade, A. *et al.* Fundamentals of microbial community resistance and resilience. *Front. Microbiol.* 3, 417 (2012).

### 71. Waring, B., Gee, A., Liang, G. & Adkins, S. A quantitative analysis of microbial community structure-function relationships in plant litter decay. *iScience* 25, 104523 (2022).

### 72. Nemergut, D. R. *et al.* Patterns and processes of microbial community assembly. *Microbiol. Mol. Biol. Rev.* 77, 342–356 (2013).

### 73. Scheel, M. *et al.* Abrupt permafrost thaw triggers activity of copiotrophs and microbiome predators. *FEMS Microbiol. Ecol.* 99, (2023).

### 74. Weissman, J. L. *et al.* Growth optimization predicts microbial success in a permafrost thaw experiment. *bioRxiv* (2025) doi:10.1101/2025.09.01.673550.

### 75. Adamczyk, M., Perez-Mon, C., Gunz, S. & Frey, B. Strong shifts in microbial community structure are associated with increased litter input rather than temperature in High Arctic soils. *Soil Biol. Biochem.* 151, 108054 (2020).

### 76. Kanevskiy, M. *et al.* Yedoma Cryostratigraphy of Recently Excavated Sections of the CRREL Permafrost Tunnel Near Fairbanks, Alaska. *Front Earth Sci. Chin.* 9, (2022).

### 77. Sellmann, P. V. *Geology of the USA CRREL Permafrost Tunnel, Fairbanks, Alaska*. https://apps.dtic.mil/sti/tr/pdf/AD0660310.pdf (1967).

### 78. Matthews, J. V., Jr. Wisconsin environment of interior Alaska: Pollen and macrofossil analysis of a 27 meter core from the Isabella basin (Fairbanks, Alaska). *Can. J. Earth Sci.* 11, 828–841 (1974).

### 79. Douglas, T. A. *et al.* Recent degradation of interior Alaska permafrost mapped with ground surveys, geophysics, deep drilling, and repeat airborne lidar. *The Cryosphere* 15, 3555–3575 (2021).

### 80. Farquharson, L. M., Romanovsky, V. E., Kholodov, A. & Nicolsky, D. Sub-aerial talik formation observed across the discontinuous permafrost zone of Alaska. *Nat. Geosci.* 15, 475–481 (2022).

### 81. Douglas, T. A., Jorgenson, M. T., Sullivan, T. & Zhang, C. Comparing thaw probing, electrical resistivity tomography, and airborne lidar to quantify lateral and vertical thaw in rapidly degrading boreal permafrost. *Cryosphere* 19, 3991–4009 (2025).

### 82. Johnson, J. B. & Lorenz, R. D. Thermophysical properties of Alaskan loess: An analog material for the Martian polar layered terrain? *Geophysical Research Letters* vol. 27 2769–2772 Preprint at https://doi.org/10.1029/1999gl011077 (2000).

### 83. Hamilton, T. D., Craig, J. L. & Sellmann, P. V. The Fox permafrost tunnel: A late Quaternary geologic record in central Alaska. *Geol. Soc. Am. Bull.* 100, 948–969 (1988).

### 84. Douglas, T. A. & Mellon, M. T. Sublimation of terrestrial permafrost and the implications for ice-loss processes on Mars. *Nat. Commun.* 10, 1716 (2019).

### 85. Burkert, A., Douglas, T. A., Waldrop, M. P. & Mackelprang, R. Changes in the Active, Dead, and Dormant Microbial Community Structure across a Pleistocene Permafrost Chronosequence. *Appl. Environ. Microbiol.* 85, e02646-18 (2019).

### 86. Amalfitano, S. & Fazi, S. Recovery and quantification of bacterial cells associated with streambed sediments. *J. Microbiol. Methods* 75, 237–243 (2008).

### 87. Morono, Y., Terada, T., Kallmeyer, J. & Inagaki, F. An improved cell separation technique for marine subsurface sediments: applications for high-throughput analysis using flow cytometry and cell sorting. *Environ. Microbiol.* 15, 2841–2849 (2013).

### 88. Poté, J., Bravo, A. G., Mavingui, P., Ariztegui, D. & Wildi, W. Evaluation of quantitative recovery of bacterial cells and DNA from different lake sediments by Nycodenz density gradient centrifugation. *Ecol. Indic.* 10, 234–240 (2010).

### 89. Berns, A. E. *et al.* Effect of gamma-sterilization and autoclaving on soil organic matter structure as studied by solid state NMR, UV and fluorescence spectroscopy. *Eur. J. Soil Sci.* 59, 540–550 (2008).

### 90. Girard-Perier, N. *et al.* Effects of X-rays, electron beam, and gamma irradiation on chemical and physical properties of EVA multilayer films. *Front. Chem.* 10, 888285 (2022).

### 91. Dittmar, T., Koch, B., Hertkorn, N. & Kattner, G. A simple and efficient method for the solid‐phase extraction of dissolved organic matter (SPE‐DOM) from seawater. *Limnol. Oceanogr. Methods* 6, 230–235 (2008).

### 92. Starr, S. F. *et al.* Organic matter composition versus microbial source: Controls on carbon loss from fen wetland and permafrost soils. *J. Geophys. Res. Biogeosci.* 130, e2024JG008445 (2025).

### 93. Hendrickson, C. L. *et al.* 21 Tesla Fourier transform ion cyclotron resonance mass spectrometer: A national resource for ultrahigh resolution mass analysis. *J. Am. Soc. Mass Spectrom.* 26, 1626–1632 (2015).

### 94. Smith, D. F., Podgorski, D. C., Rodgers, R. P., Blakney, G. T. & Hendrickson, C. L. 21 Tesla FT-ICR mass spectrometer for ultrahigh-resolution analysis of complex organic mixtures. *Anal. Chem.* 90, 2041–2047 (2018).

### 95. Kendrick, E. A mass scale based on CH2 = 14.0000 for high resolution mass spectrometry of organic compounds. *Anal. Chem.* 35, 2146–2154 (1963).

### 96. Xian, F., Hendrickson, C. L., Blakney, G. T., Beu, S. C. & Marshall, A. G. Automated broadband phase correction of Fourier transform ion cyclotron resonance mass spectra. *Anal. Chem.* 82, 8807–8812 (2010).

### 97. Savory, J. J. *et al.* Parts-per-billion Fourier transform ion cyclotron resonance mass measurement accuracy with a ‘walking’ calibration equation. *Anal. Chem.* 83, 1732–1736 (2011).

### 98. Corilo, Y. *PetroOrg Software*. (Florida State University, Tallahassee, FL, 2015).

### 99. Koch, B. P. & Dittmar, T. From mass to structure: an aromaticity index for high‐resolution mass data of natural organic matter. *Rapid Commun. Mass Spectrom.* 20, 926–932 (2006).

### 100. LaRowe, D. E. & Van Cappellen, P. Degradation of natural organic matter: A thermodynamic analysis. *Geochim. Cosmochim. Acta* 75, 2030–2042 (2011).

### 101. Liang, R. *et al.* Predominance of Anaerobic, Spore-Forming Bacteria in Metabolically Active Microbial Communities from Ancient Siberian Permafrost. *Appl. Environ. Microbiol.* 85, e00560-19 (2019).

### 102. Liang, R. *et al.* Genomic reconstruction of fossil and living microorganisms in ancient Siberian permafrost. *Microbiome* 9, 110 (2021).

### 103. Parada, A. E., Needham, D. M. & Fuhrman, J. A. Every base matters: assessing small subunit rRNA primers for marine microbiomes with mock communities, time series and global field samples. *Environ. Microbiol.* 18, 1403–1414 (2016).

### 104. Thompson, L. R. *et al.* A communal catalogue reveals Earth’s multiscale microbial diversity. *Nature* 551, 457–463 (2017).

### 105. Taylor, D. L. *et al.* Accurate estimation of fungal diversity and abundance through improved lineage-specific primers optimized for Illumina amplicon sequencing. *Appl. Environ. Microbiol.* 82, 7217–7226 (2016).

### 106. Bolyen, E. *et al.* Reproducible, interactive, scalable and extensible microbiome data science using QIIME 2. *Nat. Biotechnol.* 37, 852–857 (2019).

### 107. Callahan, B. J. *et al.* DADA2: High-resolution sample inference from Illumina amplicon data. *Nat. Methods* 13, 581–583 (2016).

### 108. Quast, C. *et al.* The SILVA ribosomal RNA gene database project: improved data processing and web-based tools. *Nucleic Acids Res.* 41, D590-6 (2013).

### 109. Robeson, M. S., 2nd *et al.* RESCRIPt: Reproducible sequence taxonomy reference database management. *PLoS Comput. Biol.* 17, e1009581 (2021).

### 110. Abarenkov, K. *et al.* The UNITE database for molecular identification and taxonomic communication of fungi and other eukaryotes: sequences, taxa and classifications reconsidered. *Nucleic Acids Res.* 52, D791–D797 (2024).

### 111. R Core Team. *R: A Language and Environment for Statistical Computing*. https://www.R-project.org/ (2021).

### 112. Whitham, J. Impact of BBDuk metagenomic read trimming and decontamination. North Carolina State Univ., Raleigh, NC (United States) https://doi.org/10.25982/77705.1341/1779218 (2021).

### 113. Bushnell, B., Rood, J. & Singer, E. BBMerge - Accurate paired shotgun read merging via overlap. *PLoS One* 12, e0185056 (2017).

### 114. Buchfink, B., Reuter, K. & Drost, H.-G. Sensitive protein alignments at tree-of-life scale using DIAMOND. *Nat. Methods* 18, 366–368 (2021).

### 115. Zhang, H. *et al.* dbCAN2: a meta server for automated carbohydrate-active enzyme annotation. *Nucleic Acids Res.* 46, W95–W101 (2018).

### 116. Kanehisa, M., Furumichi, M., Sato, Y., Matsuura, Y. & Ishiguro-Watanabe, M. KEGG: biological systems database as a model of the real world. *Nucleic Acids Res.* 53, D672–D677 (2025).

### 117. Mackelprang, R., Vaishampayan, P. & Fisher, K. Adaptation to environmental extremes structures functional traits in biological soil crust and hypolithic microbial communities. *mSystems* 7, e0141921 (2022).

### 118. McMurdie, P. J. & Holmes, S. phyloseq: an R package for reproducible interactive analysis and graphics of microbiome census data. *PLoS One* 8, e61217 (2013).

### 119. Ben-Shachar, M., Lüdecke, D. & Makowski, D. Effectsize: Estimation of effect size indices and standardized parameters. *J. Open Source Softw.* 5, 2815 (2020).

### 120. Lenth, R. V. *Emmeans: Estimated Marginal Means, Aka Least-Squares Means*. (2025).

### 121. Oksanen, J. *et al.* The vegan package. *Community ecology package* 10, 719 (2007).

### 122. Lê, S., Josse, J. & Husson, F. FactoMineR: AnRPackage for Multivariate Analysis. *J. Stat. Softw.* 25, 1–18 (2008).

### 123. Smilde, A. K., Kiers, H. A. L., Bijlsma, S., Rubingh, C. M. & van Erk, M. J. Matrix correlations for high-dimensional data: the modified RV-coefficient. *Bioinformatics* 25, 401–405 (2009).

### 124. Indahl, U. G., Næs, T. & Liland, K. H. A similarity index for comparing coupled matrices. *J. Chemom.* 32, e3049 (2018).

### 125. Rohart, F., Gautier, B., Singh, A. & Lê Cao, K.-A. mixOmics: An R package for ’omics feature selection and multiple data integration. *PLoS Comput. Biol.* 13, e1005752 (2017).

### 126. Tenenhaus, M., Tenenhaus, A. & Groenen, P. J. F. Regularized generalized canonical correlation analysis: A framework for sequential multiblock component methods. *Psychometrika* 82, 737–777 (2017).

### 127. Csárdi, G. *et al.* *Igraph for R: R Interface of the Igraph Library for Graph Theory and Network Analysis*. (Zenodo, 2025). doi:10.5281/ZENODO.7682609.

### 128. Bastian, M., Heymann, S. & Jacomy, M. Gephi: An open source software for exploring and manipulating networks. *Proceedings of the International AAAI Conference on Web and Social Media* 3, 361–362 (2009).

### 129. Anderson, M. J. Distance-based tests for homogeneity of multivariate dispersions. *Biometrics* 62, 245–253 (2006).

### 130. Stegen, J. C., Lin, X., Konopka, A. E. & Fredrickson, J. K. Stochastic and deterministic assembly processes in subsurface microbial communities. *ISME J.* 6, 1653–1664 (2012).

### 131. Sloan, W. T. *et al.* Quantifying the roles of immigration and chance in shaping prokaryote community structure. *Environ. Microbiol.* 8, 732–740 (2006).

### 132. Kembel, S. W. *et al.* Picante: R tools for integrating phylogenies and ecology. *Bioinformatics* 26, 1463–1464 (2010).

### 133. Burns, A. R. *et al.* Contribution of neutral processes to the assembly of gut microbial communities in the zebrafish over host development. *ISME J.* 10, 655–664 (2016).

### 134. Love, M. I., Huber, W. & Anders, S. Moderated estimation of fold change and dispersion for RNA-seq data with DESeq2. *Genome Biol.* 15, 550 (2014).

### 135. Kolde, R. *Pheatmap: Pretty Heatmaps*. (2019).

### 136. Cohen, J. *Statistical Power Analysis for the Behavioral Sciences*. (Routledge, London, England, 2013). doi:10.4324/9780203771587.
